# Genetic-to-Chemical Perturbation Transfer Learning Through Unified Multimodal Molecular Representations

**DOI:** 10.1101/2025.02.02.635055

**Authors:** Yiming Li, Min Zeng, Jun Zhu, Linjing Liu, Fang Wang, Longkai Huang, Fan Yang, Min Li, Jianhua Yao

## Abstract

Artificial Intelligence virtual cell (AIVC) holds transformative potential for biomedical research. Central to this vision is the systematic modeling of genetic and chemical perturbation phenotypes to accurately predict cellular dynamic states from diverse interventions. However, disparities in screening agents, library scales, experimental technologies, and data production efficiency hinder the integration, modeling, and analysis of the cross-data. Here we present **UniPert-G2CP**, a two-phase deep learning approach comprising i) UniPert, a multimodal molecular representation model that bridges genetic and chemical domains, and ii) G2CP (*Genetic-to-Chemical* Perturbation transfer learning), which systematically transforms CRISPR screen-based genetic insights into chemical perturbation modeling for cost-effective *in silico* drug screening. UniPert not only encodes multimodal perturbagens into a unified functionally interpretable sematic embedding space, but also improves phenotypic effect prediction for previously unseen gene perturbations and drug treatments. Building upon UniPert, G2CP successfully modeled large-scale cellular post-perturbation states spanning 4,994 gene and 7,821 compound perturbagens, while reducing modeling data costs by over 60%. We demonstrate that UniPert-G2CP enables efficient, generalizable simulations of multicellular, multi-domain perturbation *cause-effect* spaces, revealing differential cellular biological causality and informing mechanism-driven therapy. UniPert-G2CP opens new avenues for biological causal foundation model building, AIVC creation, and AI-powered precision medicine.

## Introduction

Simulating cellular states and predicting phenotypic effects from diverse interventions— including genetic and chemical perturbagens—has emerged as a central objective in the development of AI virtual cells^1–3^. This advancement has been fueled by the context-specific, high-throughput, multi-domain post-perturbation phenotypic profiles from initiatives such as the Connectivity Map^4,5^, Cancer Dependency Map^6,7^, and Cell Painting Gallery^8,9^. Joint modeling and analysis of cellular responses and behaviors from diverse perturbagens allows systematic exploration of biological mechanisms and therapeutic interventions, while promoting the identification of potential therapeutic targets and the discovery of novel drug candidates for specific diseases, leading to efficient data-driven personalized medicine^10–13^. This promising paradigm, however, is suffering from several critical bottlenecks in cellular perturbation modeling.

First, differences in library scales and experimental technologies between high-content genetic and chemical screens lead to a significant gap in screening space coverage and modeling data accessibility. The genetic screens with genome-scale libraries (or smaller libraries targeting ∼4,500 druggable genes^14,15^) can be performed in a pooled format using advanced CRISPR technology^16–20^, where a single cellular assay can perturb and profile various genes in bulk. In contrast, the chemical screens with enormous drug-like chemical space (estimated beyond 10^63^ compounds)^21–23^ remain physical-separated and scale-limited treatments^24,25^. Notably, even existing largest chemical perturbation atlases only cover thousands small molecules^5,26^, while full-scope screening is experimentally infeasible and necessitate computational simulation. These gaps exacerbate the imbalance in data productivity and modeling needs between the two domains.

Second, the inherent molecular-modal disparities between perturbagens constrain their universal representation and further limit cross-domain perturbation connectivity and modeling. Advanced encoders for large biomolecules and small molecules typically accept input of nucleotide/amino-acid-scale biosequences^27–29^ and atomic-scale SMILES^30–32^, respectively. Numericizing these molecules (with varying scales and lengths) into a shared interpretable space i) requires rich empirical knowledge to establish their biological relations, and ii) risks losing crucial local information of the molecules, such as evolutionary conserved regions of biosequences^33^ and functional groups of compounds^34^.

Third, modeling towards high cellular heterogeneity and generalization constitutes a grand challenge^35–37^. The utility of perturbation phenotypic data can be characterized along two dimensions: depth, referring to the sample size (i.e., number of post-perturbation profiles) per experimental condition; and breadth, denoting the diversity of conditions (i.e., perturbagen counts). While single-cell technologies have substantially increased data depth ^38^, these benefits are largely confined to interpolation within observed perturbation distributions, with limited gain in predicting novel, out-of-distribution (OOD) conditions. Previous studies have shown that cross-context transfer learning can improve model generalization in new cellular environments^39,40^,but it often tends to be overwhelmed by common patterns and obscures context-specific signals, thereby compromising cellular heterogeneity learning.

Training efficient multicellular perturbation effect prediction models requires experimental data with sufficient perturbation breadth across diverse cellular contexts. However, conducting chemical screening with sufficient coverage remains impractical due to its vast chemical space and labor-intensive experimentation^41^. Given that chemical perturbations can mimic genetic effects by converging on shared signaling cascades and regulatory programs^42–47^, a promising strategy is to pre-train models using genetic screen profiles (with more systematic yet smaller libraries) and fine-tune them with limited chemical screen profiles. This strategy leverages pre-learned unique genetic background information to enable cost-efficient, context-aware chemical perturbation modeling^46^ as well as generalizable phenotypic predictions for numerous untreated compounds, further supporting the systematic parsing of compound mechanisms of action (MOA) and cellular response sensitivity, and propelling the evolution from chemical genetics to chemical genomics^48,49^.

In this study, we developed a two-phase deep learning approach to overcome modeling bottlenecks in phenotypic screening across perturbation domains, molecular modalities, and library scales. In phase 1, we developed UniPert, a contrastive learning-based^50^ multimodal molecular representation learning model that encodes both large biomolecules and small molecules into a shared sematic embedding space, bridging the genetic and chemical perturbation domains. In phase 2, we integrated UniPert into perturbation effect prediction model to develop G2CP, a *genetic-to-chemical* perturbation transfer learning framework that leverages systematic CRISPR-based genetic screening to boost *in silico* drug screening efficiency.

We proceed to provide an overview of UniPert’s architecture (Figure 1a) and demonstrate its versatility to i) generate multimodal molecule embeddings with significant biological and pharmacological interpretability, and ii) be plugged in advanced perturbation predictors and improve their performance in dealing with unseen perturbagens across various perturbation domains, complexities and data scales (Figure 1e, left and middle). Then, we present the overview of G2CP (Figure 1b) and evaluate it on transcriptomic and morphological profiles containing dual-domain perturbations across multiple cellular contexts, demonstrating its superiority in cost-efficient *in silico* drug screening (Figure 1d). Last, we utilize UniPert-G2CP to generate perturbation *cause-effect* representation spaces for 4,994 genes and 7,821 small molecules across five cancer cell lines (Figure 1c), and perform a joint analysis of these spaces to character the drug perturbation cellular specificity (Figure 1e, right).

**Figure 1.**
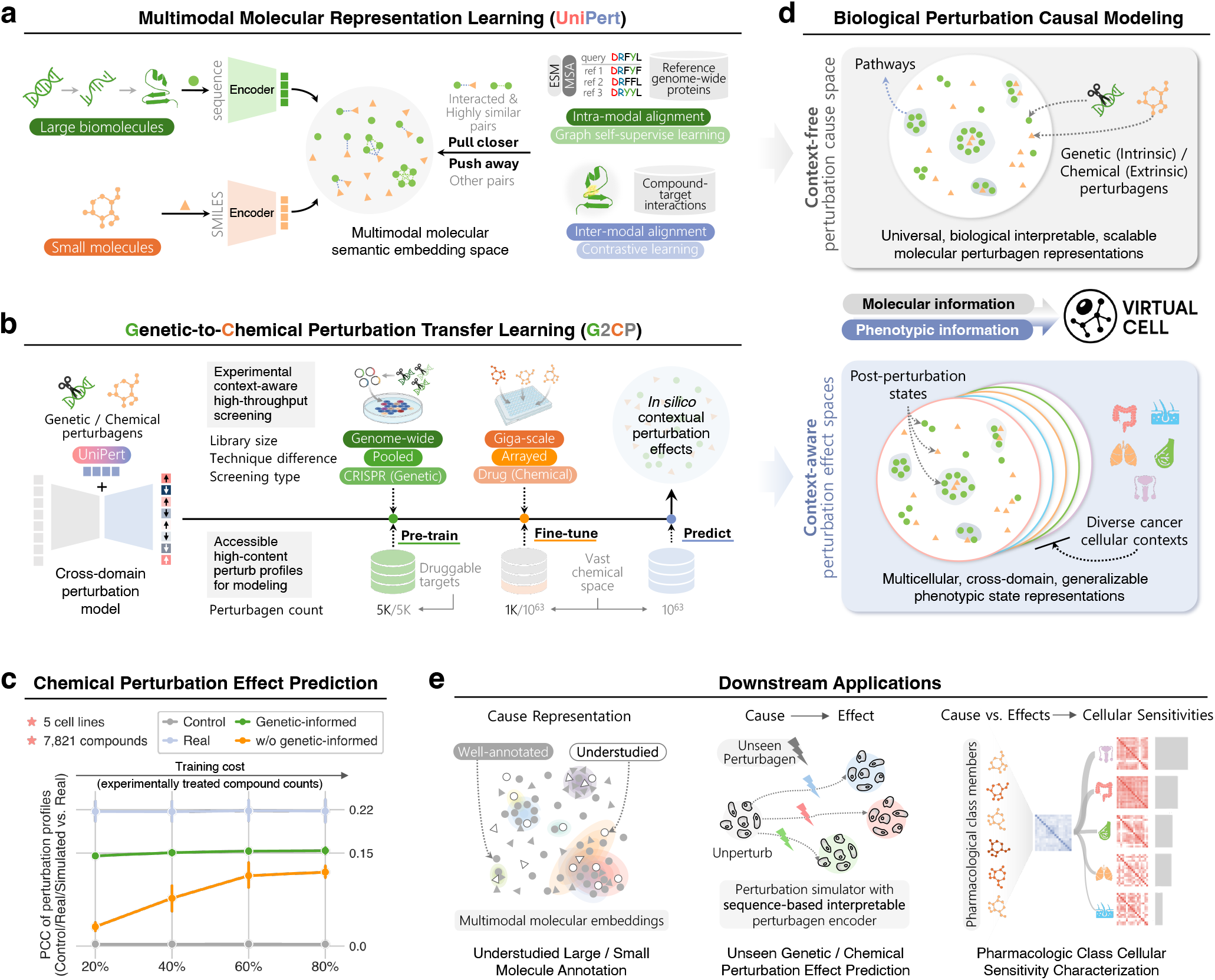
Overview of UniPert-G2CP. **(a)** Illustration of UniPert model. UniPert learns interpretable multimodal molecular representations by integrating computation techniques (right)—including protein language model (ESM), multiple sequence alignment (MSA), graph self-supervised learning, and contrastive learning—and leveraging priori experimental data of compound-target interactions, connecting the underlying multimodal molecular intervention factors (perturbagens) between genetic and chemical perturbation domains. UniPert owns versatile and scalable encoders (left) for both large biomolecules (green circles) and small molecules (orange triangles), which pulls closer (dotted lines between molecular subjects) interacting compound and target pairs and highly similarity target pairs, while pushing away other functionally unrelated pairs (circle/triangle pairs without lines) for the multimodal subjects within the sematic embedding space (middle). **(b)** Illustration of G2CP framework. In large-scale cell-based screening, chemical perturbations are more lab-costly than genetic perturbations due to their different screen library scales and techniques (top). Thus, G2CP employs a transfer learning framework with “genetic pre-train + chemical finetune” strategy for cost-efficient chemical perturbation modeling and prediction to the vast chemical space (bottom). “Genetic pre-train” enables models to pre-learn systematic contextual genetic backgrounds and informs chemical “fine-tune”. This cross-domain modeling strategy is benefited from the unified perturbagen embedding space constructed by UniPert (left). **(c)** Overall evaluation of the genetic-informed perturbation models for unseen chemical perturbation effect prediction involving 7,821 compounds across five cancer cell lines. “Genetic-informed” means models trained using G2CP strategy, which reducing experimental chemical screen costs by over 60%. **(d)** UniPert and G2CP together achieve generalizable biological causal modeling by efficiently representing context-free molecules (top) and context-aware phenotypes (bottom), which are fundamental for AI virtual cell building (middle). **(e)** UniPert-G2CP supports a series of downstream applications: encoding and annotating understudied large biomolecules and small molecules with interpretable embeddings (left), ii) improving perturbation effect prediction for previously unseen genes and compounds (middle), and iii) uncovering cellular sensitivities of pharmacological classes through joint analysis of simulated large-scale perturbation cause and effect spaces across multiple cellular contexts (right).

## Results

### UniPert Bridges Genetic and Chemical Screen Domains by Learning a Unified Perturbagen Representation Space

Genetic and chemical perturbation screen studies remain largely disconnected due to the distinct perturbagens, especially for those unannotated. To bridge these two domains, we propose to develop a multimodal molecular representation model (UniPert) that leverages priori biological information to unify multi-domain perturbagens and facilitate cross-domain phenotypic connections (Extended Data Figure 1a). The goal of UniPert is to encode diverse perturbagens into a shared embedding space from their original sequences, aggregating i) genetic perturbagens with similar (or close) functional roles, ii) chemical perturbagens with the same MOA, and iii) genetic and chemical perturbagens involved in the same biological pathway. To achieve this goal, UniPert is designed with multiscale molecular inputs, adapted model encoders, and combined training strategies (Extended Data Figure 1b).

For chemical perturbagens, UniPert accepts SMILES as inputs, ensuring the encoder’s scalability for the vast chemical space. The SMILES strings are initially embedded by binary topological extended-connectivity fingerprints (ECFPs)^30^ for substructure characterization. These sparse features are then converted into dense embeddings for refining the essential functional groups and enabling alignment with genetic perturbagen embeddings.

For genetic perturbagens, considering that proteins are the fundamental units of cellular function and central roles in both screen domains, UniPert encodes target genes by translating them into amino acid sequences. Unlike small molecules, proteins own longer sequences and more complex biological patterns. Advanced large protein language models (PLMs), such as ESM^27^, have been proved to generate informative protein embeddings and are promising for genetic perturbagen representation. However, directly compressing these high-dimensional, residue-level embeddings into global, protein-level embeddings risks losing important functional local sequence characteristics. To address this challenge, UniPert combines advanced PLM^27^ with traditional multiple sequence alignment (MSA) method^51^, leveraging graph neural networks (GNNs)^52^ to capture both global contextual features and local conserved motif patterns. Specifically, for each query protein, UniPert performs Smith-Waterman-based local alignment^53^ against a reference set of genome-wide proteins (n = 19,187), constructing a query-centered weighed graph in which nodes represent proteins and edges encode pairwise sequence similarities. Initial node embeddings are generated from ESM^27^ and subsequently refined through GNN-based message passing mechanism, enabling functional information propagation among similar sequences. This hybrid strategy enhances UniPert’s ability to identify conserved sequence regions and capture protein functional similarities.

UniPert adopts a dual optimization strategy to establish both intra-modal and inter-modal relationships within a shared latent semantic embedding space (Extended Data Figure 1b, right). On one hand, a graph self-supervised learning strategy^54^ (Methods) was employed to complement the sequence alignment algorithm and the PLM-based representations in learning robust protein embeddings. This strategy encourages consistency in protein node embeddings across different graph variants generated by randomly dropping edges and node features. On the other hand, UniPert maps compounds and proteins in the embedding space by integrating a contrastive learning-based alignment strategy^55^ (Methods) with a large collection of compound-target interactions, including 81,397 pairs involving 4,250 proteins and 8,533 compounds. This inter-modal alignment adjusts distances between compound– target pairs, drawing interacted molecules closer in the embedding space and capturing their shared roles in biological pathways.

### UniPert Generates Functional-Informed Embeddings and Facilitates Annotation for Understudied Molecules

Annotation inequality among molecules hinders biomedical progress, as most studies focuses on a limited set of increasingly well-knowns^56^. The Understudied Proteins Initiative seeks to bridge this gap by systematically associating uncharacterized proteins with annotated counterparts^57,58^. Such associations can be fulfilled by integrating similarity-based approaches with interpretable molecular sematic representations^59,60^. To this end, we assess the functional and biological interpretability of UniPert-generated embeddings and their capacity to support annotation for small molecules and large biomolecules across diverse taxonomy systems, such as drug MOA classes, protein pharmacological classes, protein family classes and hierarchies (Figure 2a).

**Figure 2.**
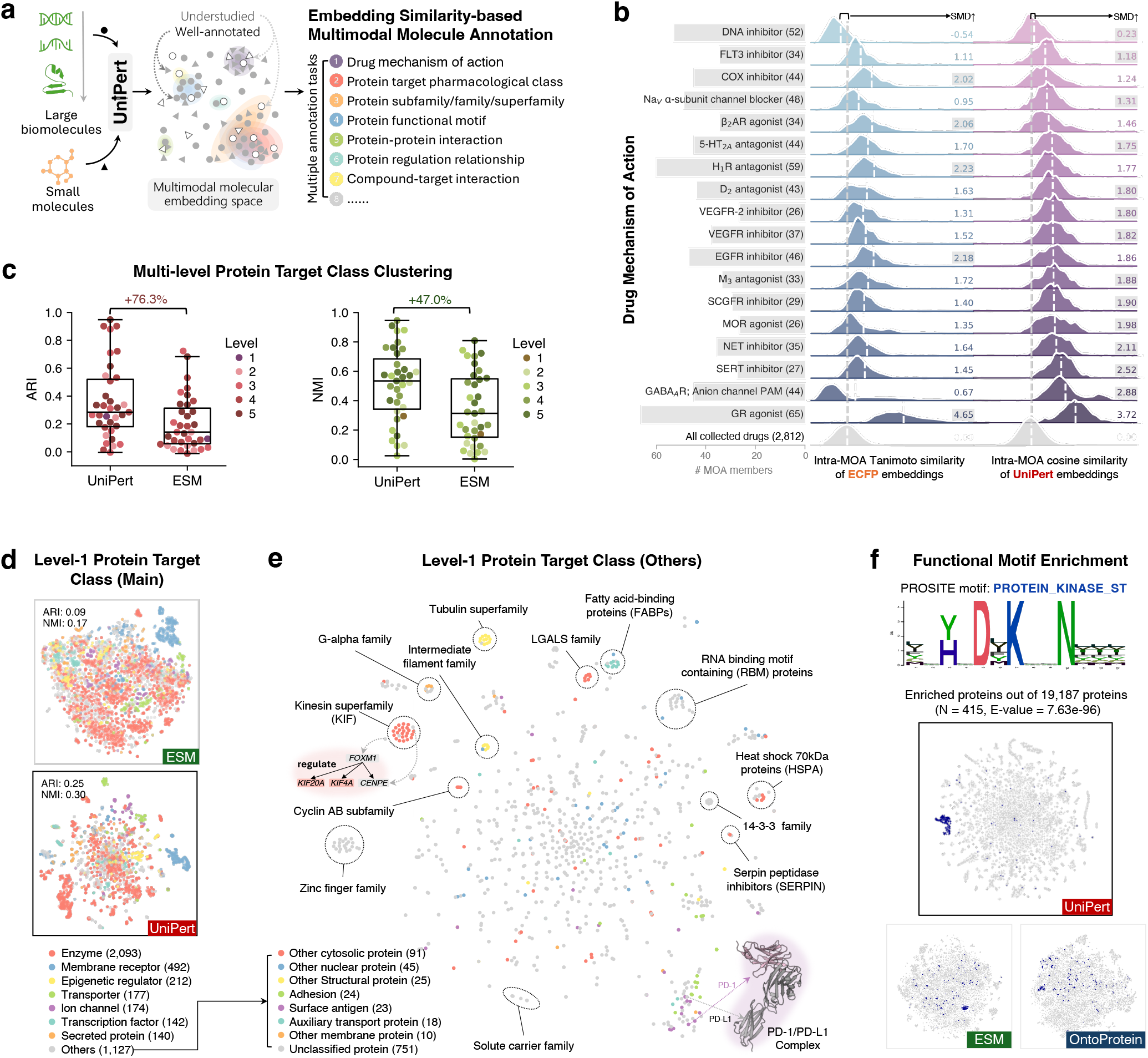
UniPert generates interpretable compound and protein embeddings and facilitates the annotation of understudied molecules. **(a)** UniPert generates interpretable embeddings for both large biomolecules (circles) and small molecules (triangles) within a unified molecular embedding space (left). These semantic embeddings support diverse class annotation tasks for understudied/poor-annotated molecules (white-filled circles/triangles), including drug mechanism of action (MOA) classification, protein target pharmacological classification, protein subfamily/family/superfamily classification, protein functional motif annotation, protein-protein interaction annotation, protein regulatory relationship annotation, and compound-target interaction prediction (right). **(b)** Semantic similarity analysis of UniPert and ECFP embeddings across MOA-defined drug classes. Each row represents the pairwise drug embedding similarity distribution among drugs within a given MOA class (> 25 members), measured using cosine similarity (UniPert) or Tanimoto similarity (ECFP). The standardized mean difference metric, SMD, was employed to quantify the semantic separability between drugs within each MOA class and the overall drug population (n = 2,812). Higher SMD values (marked with a grey background) indicate greater semantic distinctiveness of a given MOA class in the embedding space. **(c)** Clustering performance comparison of UniPert and ESM embeddings across 35 protein target pharmacological classes spanning 5 hierarchical levels, with class hierarchical relationships illustrated in Extended Data Figure 2a. Boxplots show Adjusted Rand Index (ARI) and Normalized Mutual Information (NMI) scores across multiple class levels; each data points correspond to a specific class. **(d)** t-SNE visualization and clustering comparison of 7 major level-1 target classes (legend as bottom) using UniPert (middle) and ESM (top) embeddings. **(e)** t-SNE visualization of UniPert embeddings for proteins belonging to the remaining seven minority classes and the unclassified group with the legend at left bottom. Clustering of protein families is evident. FOXM1 proximity to kinesin superfamily proteins reflects their regulatory relationships (left). PD-1/PD-L1 subunit proximity highlights their interaction (bottom right). **(f)** Functional motif enrichment analysis. Sequence logo (top) illustrates the most significant PROSITE motif, i.e., PROTEIN_KINASE_ST (E-value = 7.63^−96^), identified in 415 out of 19,187 proteins using the MEME suite. t-SNE plots (bottom) compares UniPert, ESM, and OntoProtein embeddings of the enriched proteins.

For small molecule drugs, we compiled 2,812 small molecule drugs with known MOAs and performed semantic similarity analysis on them (Supplementary Table 1). We examined embedding similarities for all drug pairs and compared the similarity distributions between drug pairs within the same MOA category and all drug pairs (overall group). Compared to ECFP^30^, UniPert-generated drug embeddings demonstrate improved inter-MOA similarity separation in 13 out of 18 MOA categories (Figure 2b). Intra-MOA members exhibit higher embedding similarity than those with distinct MOAs, demonstrating UniPert’s ability to encode mechanistic relationships. Moreover, we observed that drug pairs within two MOA categories—’DNA inhibitor’ and ‘GABAAR; Anion channel PAM’—exhibited significantly low similarity based on ECFP embeddings, suggesting that the traditional fingerprints-based representation method fails to capture the shared mechanism relationships in these categories. In contrast, UniPert brings them closer in the semantic representation space, thereby enhancing the potential for discovering novel drugs within these challenging MOA categories.

For protein targets, we first performed clustering analysis for 4,417 human protein targets with expert-curated hierarchical pharmacological classifications (Supplementary Table 2). These classifications span from broad target class to more specific subclasses such as functional families and binding domains, reflecting the increasing granularity in biological and pharmacological relationships. Compared to ESM, UniPert more accurately clusters 35 target pharmacological classes (PCLs) across multiple hierarchical levels, achieving increases of 76.3% in Adjusted Rand Index (ARI) and 47.0% in Normalized Mutual Information (NMI), respectively (Figure 2c). In-depth embedding visualization and clustering comparisons across 5 major target classes (enzyme, membrane receptor, epigenetic regulator, transporter, and ion channel) and their subclasses demonstrate that UniPert learns an interpretable embedding space that spatially enriches proteins with pharmacological relationships at a fine-grained scale (Figure 2d and Extended Data Figure 2).

Further investigation of minority and unclassified targets reveals UniPert’s ability to assist in annotating protein family classification, protein regulatory relationships, and protein-protein interactions (Figure 2e). For example, UniPert successfully clusters protein families such as zinc finger proteins, the kinesin superfamily proteins (KIFs), and tubulin proteins. Notably, UniPert locates previously unclassified proteins CENPE (centrosome protein E, also known as KIF10) and FOXM1 (forkhead box protein M1, a key transcription factor regulating the expression of cell cycle genes) closing to the KIFs, consistent with the reported KIF membership of CENPE^61^ and the regulatory relationships between FOXM1 and KIFs^62–64^ (Figure 2e, left). UniPert also captures the interaction between subunits of the asymmetric PD-1/PD-L1 complex^65^ (Figure 2e, bottom right). Moreover, compared to other advanced PLMs including ESM^27^ and OntoProtein^28^, UniPert presents significant advantages in organizing stratified protein family structures, i.e., superfamily, family, and subfamily (Extended Data Figure 3), and in enriching proteins with conserved functional motifs (Figure 2f and Extended Data Figure 4). UniPert’s superior motif-level characterization and enrichment capacity offers a unique advantage for uncovering conserved functional elements, enabling more precise functional inference, novel target discovery, and cross-species annotation transfer in under-characterized protein spaces.

### UniPert Enhances Effect Prediction for Unseen Single- and Multi-gene Perturbation

Biological perturbation modeling has gained increasing attention due to the growing availability of high-throughput screening data^2^. However, existing models often struggle to generalize to previously untreated experimental conditions^35^. To assess UniPert’s ability to assist in such out-of-domain settings, we systematically benchmarked it within two representative perturbation effect prediction frameworks—GEARS^66^ and CPA^67^—across a range of datasets spanning diverse perturbation domains, complexities and scales (Figure 3a).

**Figure 3.**
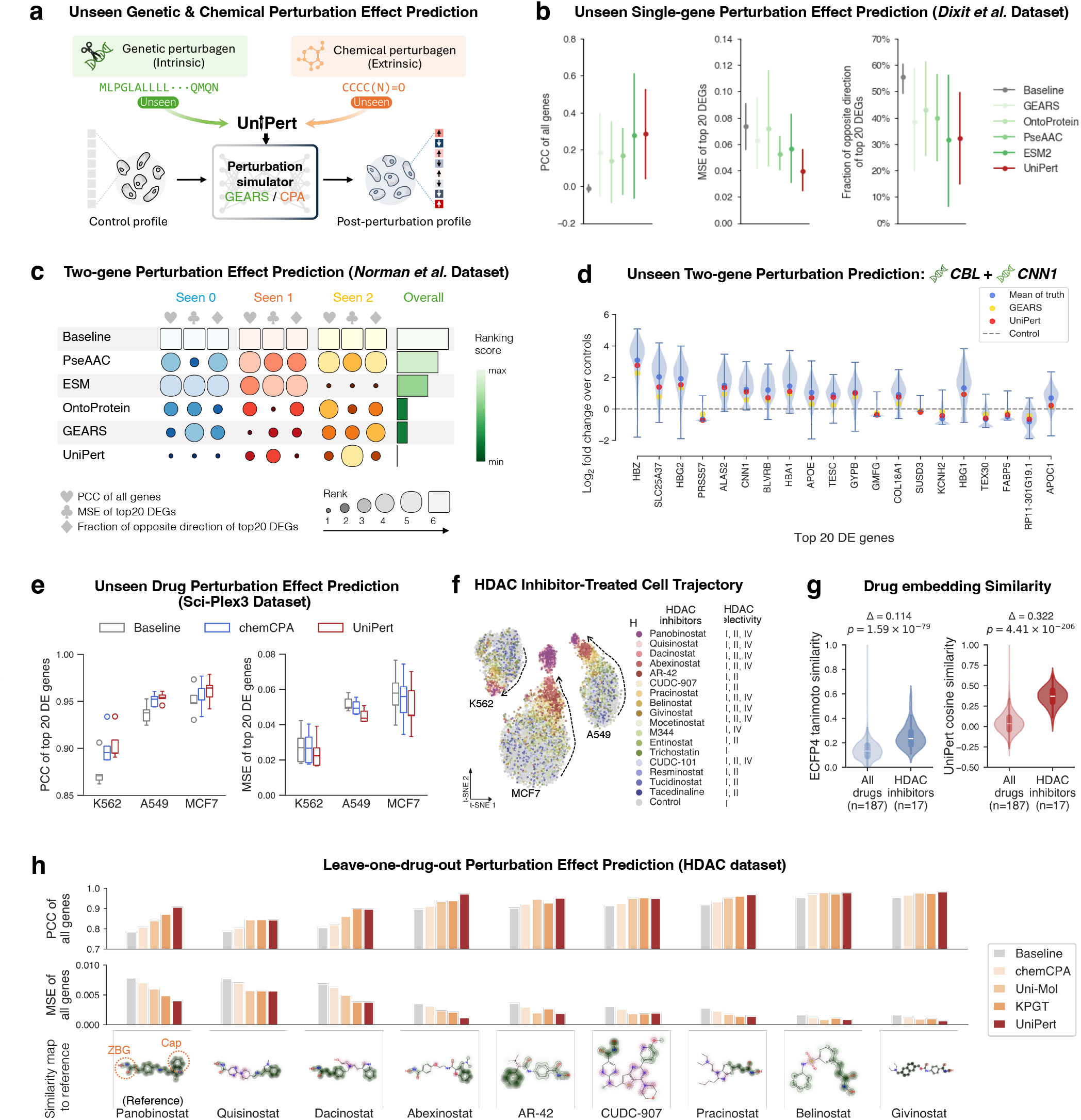
UniPert enhances unseen genetic and chemical perturbation effect prediction. **(a)** UniPert functions as a sequence-based perturbagen encoder within perturbation predictors to improve their prediction generalization and efficiency for unseen conditions. **(b)** Unseen single-gene perturbation effect prediction performance on the *Dixit et al*. dataset. Metrics shown are Pearson correlation coefficient (PCC) of all genes, mean squared error (MSE) of top 20 differentially expressed genes (DEGs), and fraction of opposite direction of top 20 DEGs. Error bars indicate 95% confidence intervals calculated from five independent trials using varying random seeds for data splitting. **(c)** Two-gene perturbation effect prediction performance on the *Norman et al*. dataset. The heatmap shows mean ranking scores from 5 trials across different evaluation metrics for three testing scenarios: 0, 1, or 2 genes seen during training in gene combinations. **(d)** Predicted changes over controls of the top 20 DEGs resulting from perturbation of the unseen gene combination (*CBL* + *CNN1*). Violin plots show gene expression change distributions of real observed post-perturbated profiles with blue dots as their mean. Red and yellow marks indicate mean predictions from UniPert and GEARS. The grey dotted line represents the mean unperturbed control expression. **(e)** Unseen drug perturbation effect prediction performance on the sci-Plex3 dataset. Boxplots show PCC and MSE for K562, A549, and MCF7 cell lines. Evaluation results are shown for different experimental trials from five data splits, with error bars representing 95% confidence intervals. Results for multiple DEG scales can be found in Extended Data Figure 6. **(f)** Cell response trajectories of HDAC inhibitor treatments. t-SNE visualization of cell states for different HDAC inhibitors. The black dotted line shows the trajectory from control (vehicle-treated) cells and towards cells exhibiting stronger responses under 1μM drug treatment. **(g)** Pairwise drug embedding similarity. Violin plots show the distribution of ECFP4 Tanimoto similarity (left) and UniPert cosine similarity (right) for the ‘All drugs’ (n = 187) and ‘HDAC inhibitors’ (n = 17) groups. Mean differences (Δ) and statistical significance (p-values) were calculated using an independent samples t-test. **(h)** Leave-one-drug-out perturbation effect prediction performance on the HDAC dataset. Top and middle panels show the PCC and MSE metrics results. The bottom panel exhibits 2D molecular structures and structural similarities between Panobinostat (reference) and other eight HDAC inhibitors, which highlights the zinc-binding group (ZBG) and Cap regions. In b), c), e) and h), the baseline represents metric calculation of real unperturbed profiles vs. real post-perturbed profiles, demonstrating the actual phenotypic differences between the control and treatment groups of the test perturbagens.

We first investigated the role of UniPert within the GEARS-derived framework (Extended Data Figure 5a, bottom). GEARS^66^ learns functional gene perturbagen embeddings by integrating Gene Ontology (GO) priors with GNNs. While effective, the reliance on curated ontologies limits its scalability and robustness, as the biological relationships learned in the embeddings depend heavily on the quality and completeness of the underlying annotations. In this regard, we extended GEARS framework by incorporating several alternative pre-trained/pre-defined protein encoders, including PseAAC^68^, ESM^27^, OntoProtein^28^ and UniPert, and compared their performance (Extended Data Figure 5a, top).

For single-gene perturbations, we evaluated the prediction performance on the benchmark Perturb-seq dataset from *Dixit et al*.^17^. UniPert presents an optimal performance across three evaluation metrics compared to competing encoders (Figure 3b). In the cases of perturbing unseen cell-cycle regulator *CEP55* and transcription factors *AURKA* and *AURKB*, UniPert showed higher correlation between predicted and observed gene expression changes compared to the original GEARS model (Extended Data Figure 5b).

We next evaluated UniPert on the more complex task of combination gene perturbation prediction based on the *Norman et al*. dataset, which includes three test scenarios: i) both genes in the combination were seen during training, ii) one gene was seen, and iii) neither gene was seen. Across these scenarios, UniPert outperformed other methods in the latter two, especially when both genes were unseen during model training (Figure 3c). More accurate prediction of the post-perturbation expression changes of the top 20 differential expressed genes (DEGs) highlights UniPert’s power in enhancing multigene perturbation simulation. For example, when perturbing the unseen gene combination of *CBL* and *CNN1*, UniPert’s predictions are more closely aligned with the observed than GEARS (Figure 3d). Moreover, UniPert also maintained consistent advantages when predicting the effect of individual combination member gene (Extended Data Figure 5c), similar improvements observed in another case involving gene combination of *FOSB* and *CEBPB* (Extended Data Figure 5d).

### UniPert Improves Effect Prediction to Unseen Drug Treatments Across Data Scales

Beyond genetic perturbations, we also evaluated UniPert for predicting cellular responses after compound treatments within the CPA-derived framework (Extended Data Figure 6a, bottom). CPA^67^ is an autoencoder-based perturbation simulator available for complex seen conditions. Its extension, chemCPA, implements the prediction for unseen compounds by incorporating an ECFP encoder. This encoder can be replaced with alternative molecular representation methods—such as Uni-Mol^31^, KPGT^32^, and our proposed UniPert (Extended Data Figure 6a, top)—to enhance prediction for OOD conditions.

We benchmarked these encoders within the CPA-derived framework using the large-scale sci-Plex3 drug screening dataset^24^ (Extended Data Figure 6b). The dataset was split hierarchically by pathway, with 20% of drugs from pathways containing over 10 members held out as unseen compounds for testing, and the rest used for training. The results show that UniPert outperformed chemCPA in terms of Pearson correlation coefficient (PCC) and mean squared error (MSE) at multiple top DEG scales across different cellular contexts (Figure 3e and Extended Data Figure 6c, d).

In addition to evaluation on the large-scale dataset, we also conducted leave-one-drug-out evaluations on a focused subset of histone deacetylase (HDAC) inhibitors. Prior studies have shown that drugs targeting epigenetic regulators often induce highly similar transcriptional responses^24,69^. Accordingly, we extracted post-perturbation gene expression profiles from the sci-Plex3 dataset following treatments with 17 HDAC inhibitors at 1µM concentration for 24 hours. The projected lower-dimensional visualization demonstrated coherent cell response trajectories across three contexts treated with these HDAC inhibitors, with Panobinostat eliciting the strongest response (Figure 3f, left). Furthermore, HDAC inhibitors with significant responses tend to be non-selective, such as Panobinostat, Quisinostat, and Abexinostat, while less potent inhibitors such as Tacedinaline displayed higher selectivity for specific HDAC targets (Figure 3f, right).

Subsequent analysis demonstrated that UniPert embeddings effectively captured the pharmacological distinctions of HDAC inhibitors, outperforming ECFP (Figure 3g). In leave-one-drug-out evaluation of the top nine most responsive HDAC inhibitors, UniPert showed superior performance in handling unseen drugs compared to other methods (Figure 3h). Structural comparisons further revealed that these potent inhibitors shared key features in the zinc-binding group (ZBG) and cap regions, in line with established pharmacological knowledge^70^, suggesting UniPert’s ability to capture meaningful molecular features.

### G2CP Enables Cost-Efficient *In Silico* High-Content Chemical Screening via Pre-Learning Genetic Contexts

Contemporary molecular targeted therapy requires a closed loop that links disease mechanism elucidation with cellular context-aware drug screening^71^. While genome-wide association studies and CRISPR-based functional genomics have greatly facilitated the mapping of genotype-phenotype relationships and the identification of individual-specific targets^72–75^, their translational efficiency in preclinical phase remains constrained by the experimental bottleneck of extensive iterative drug screening. Computer-aided high-throughput approaches such as molecular structure-based virtual screening^76,77^, can accelerate early-stage drug design; however, they face inherent limitations in simulating cellular heterogeneous of drug responses^78^ (Figure 4a). These limitations underscore the need for integrated computational frameworks that jointly incorporate molecular characters and phenotypic contexts for context-aware drug screening *in silico* ^79,80^.

**Figure 4.**
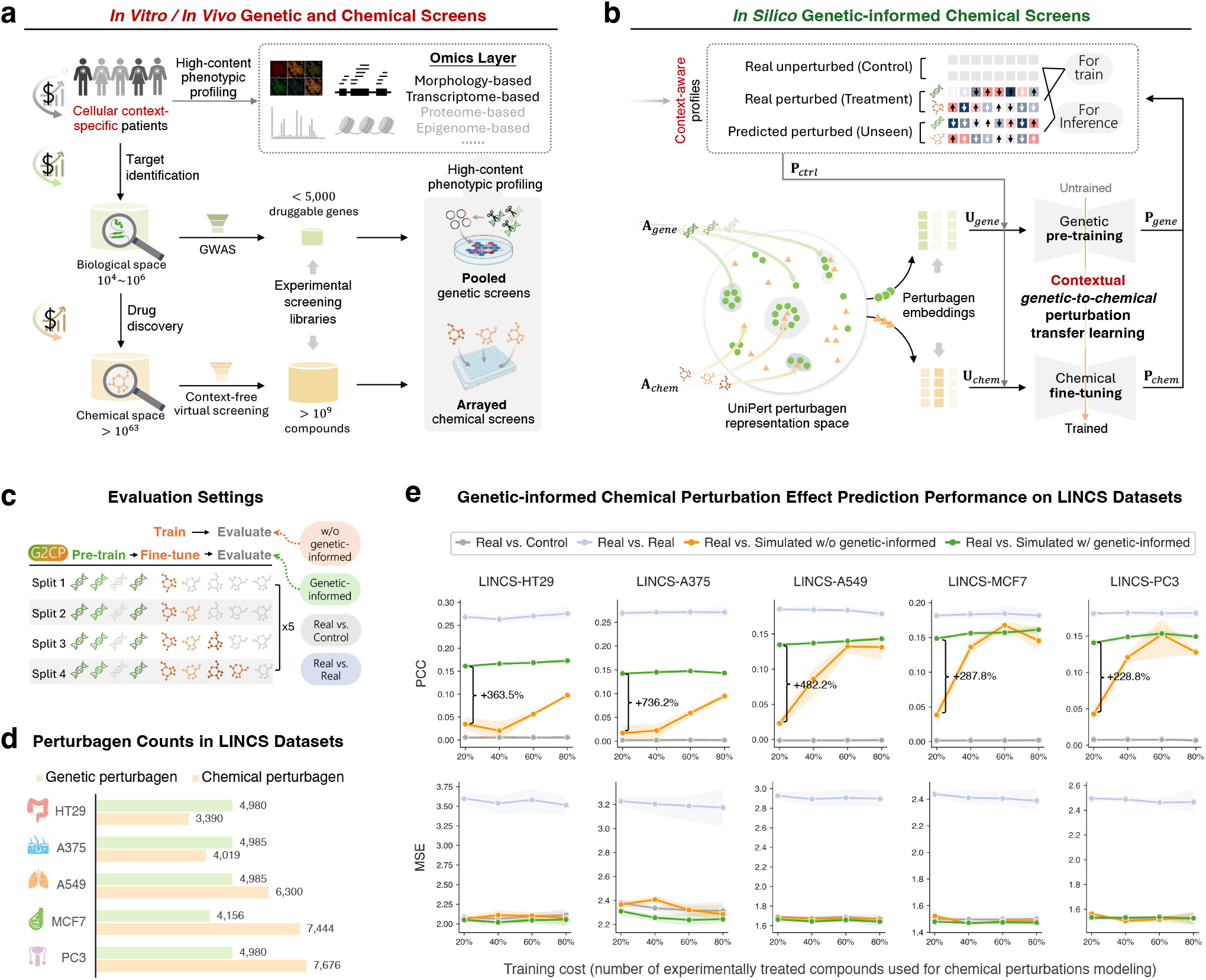
G2CP enables cost-effective, genetic-informed chemical perturbation modeling. **(a)** Schematic of the envisioned molecular targeted therapy process involving both *in vitro/in vivo* genetic and chemical perturbations. This process is resource-intensive due to the heterogeneity of cellular contexts, the vast size of chemical libraries, and the cumbersomeness and costliness of large-scale arrayed chemical screens. **(b)** Overview of the G2CP framework for computationally simulating large-scale *in vitro/in vivo* drug screening via genetic-informed chemical perturbation modeling. G2CP leverages a unified multimodal perturbagen embedding space (left) and adopts a cross-domain modeling strategy (see Methods) comprising genetic pre-training and chemical fine-tuning stages (right). P_*ctrl*_, P_*gene*_, P_*chem*),_ represent phenotypic profiles measured under control (no perturbation), genetic perturbation, and chemical perturbation conditions, respectively. A_*gene*_ and A_*chem*,_ represent the corresponding perturbagen annotations of perturbation conditions. **(c)** Evaluation settings for G2CP. Performance is assessed by varying the proportion of chemical conditions used for training/fine-tuning and the rest for OOD testing. Each data split is evaluated across five independent trials with different random seeds. **(d)** Perturbagen information in the transcriptome-based LINCS datasets. Bar plots show the number of genetic and chemical perturbagens for five cancer cell lines: HT29, A375, A549, MCF7, and PC3. **(e)** Performance of G2CP in unseen chemical perturbation effect prediction on the LINCS datasets. Point plots show performance trends (mean ± 95% confidence interval) as the number of training compounds increases. ‘Real vs. Control’ refers to the baseline setting where experimentally observed post-perturbation effects relative to the unperturbed; ‘Real vs. Real’ represents the baseline comparing the post-perturbation effects observed under identical experimentally conditions to assess inherent variability; ‘Real vs. Simulated w/o genetic-informed’ represents baseline models trained without the genetic pre-training stage; ‘Real vs. Simulated w/ genetic-informed’ represents models training using G2CP.

To this end, we present G2CP, a *genetic-to-chemical* perturbation transfer learning framework that builds on UniPert’s ability to functionally encode multimodal molecules and enhance multi-domain perturbation predictions. In G2CP, the perturbation prediction model is first pre-trained on massive genetic screen data to learn cellular genetic backgrounds and response patterns tied to distinct targeting pathways, and subsequently fine-tuned using chemical perturbation data to adapt to compound response dynamics (Figure 4b, Methods). By leveraging transfer learning and systematic pooled CRISPR screens, G2CP compensates for the sparse coverage of chemical perturbation space in experimental data for model training, facilitating cost-efficient context-aware high-content *in silico* drug screening.

To validate G2CP’s efficacy, we curated publicly available large-scale and dual-domain phenotypic screening data, including the LINCS^5^ and CPJUMP1^81^ datasets (Supplementary Table 3). The LINCS datasets comprises transcriptomic perturbation profiles (featuring 978 landmark transcripts) resulting from over ten thousand CRISPR gene knockouts and compound treatments across five different cancer cell lines (Figure 4d). The CPJUMP1 datasets contain well-level morphological profiles (with 838-dimensional extracted features) involving 160 gene and 302 drug perturbagens across two cell lines (Extended Data Figure 7a, b). We conducted a comprehensive evaluation by holding out varying proportions (20%, 40%, 60%, and 80%) of chemical perturbation conditions as OOD test sets, and compared G2CP against three baselines: i) a null model (real vs. control), ii) the consistency between replicates of the same perturbagen (real vs. real), iii) the prediction model trained under identical settings but without genetic pre-training (Figure 4c).

For the transcriptomic LINCS datasets, G2CP demonstrated remarkable improvements in terms of PCC (Fig. 4e, top). Without genetic pre-training, the PCC increased gradually as more compounds used for training. In contrast, the genetic-informed model consistently delivered significantly more robust and accurate predictions across five contexts, achieving improvements ranging from 228.8% to 736.2%. Notably, even with only 20% of fine-tuning conditions, the pre-trained models outperformed models trained from scratch using 80% of compounds, effectively reducing the experiment cost for chemical perturbation modeling by up to 60%. Moreover, we noticed that datasets with fewer chemical perturbagens (< 5000), such as the A375 and HT29 datasets, struggled to achieve desirable performance even with 80% of training data, while genetic pre-training compensated for this limitation. Additionally, unlike PCC, we observed that the MSEs between replicates of real measured post-perturbation effects induced by the same perturbagen were consistently higher than those of other baselines and G2CP predictions (Figure 4e, bottom). This indicates that gene expression changes induced by a same perturbagen are more consistent and predictable in direction than in magnitude.

For the morphological CPJUMP1 datasets, the results show that both pre-trained and non-pre-trained models exhibited limited prediction performance in two cell lines (Extended Data Figure 7c). Although genetic pre-training enhanced prediction accuracy in certain trials, these improvements are not substantial. These limited performances can be attributed to several factors: i) the limited number of perturbation conditions in both domains results in insufficient training data for capturing meaningful patterns, ii) the suboptimal morphological profile extraction method fails to adequately represent post-perturbation states, and iii) the insufficient quality control, such as potential well position effects, may introduce significant noise into the data (Extended Data Figure 7d). Similar concerns regarding these limitations have been raised in recent studies^8,82^.

### UniPert-G2CP Simulates Multicellular Perturbation Cause and Effect Representation Spaces

Having validated the effectiveness of UniPert and G2CP in modeling biological perturbation causes (perturbagens) and their resulting effects (phenotypic states) through extensive benchmarking, we further extended its application to the joint analysis of perturbation causality across multiple cellular contexts. This extension enables a systematic investigation of cellular heterogeneity in perturbation responses, which is critical step toward identifying context-specific therapeutic strategies in precision medicine^83–87^.

To rigorously construct context-specific perturbation prediction models and perform unbiased cross-context evaluation, we standardized the training pipeline across five LINCS datasets. Drawing on the original Connectivity Map (CMap) workflow^5^ of categorizing perturbagens into well-annotated (*Touchstone*) and unannotated (*Discovery*) sets, we adapted this classification scheme to G2CP-based model training. Specifically, for each cell line dataset, we designated all genetic perturbagens (∼5,000 genes) as the *Guide* set for pre-training, 1,012 chemical perturbagens (overlapping with the CMap *Touchstone* and involved in 452 pathways) as a refined *Touchstone* set for fine-tuning, and the remaining 6,800 compounds (ranging from 2,378 to 6,664 per dataset) as refined *Discovery* sets for evaluation. This procedure resulted in five independently trained, context-specific perturbation predictors under harmonized G2CP training protocols (Figure 5a, left).

**Figure 5.**
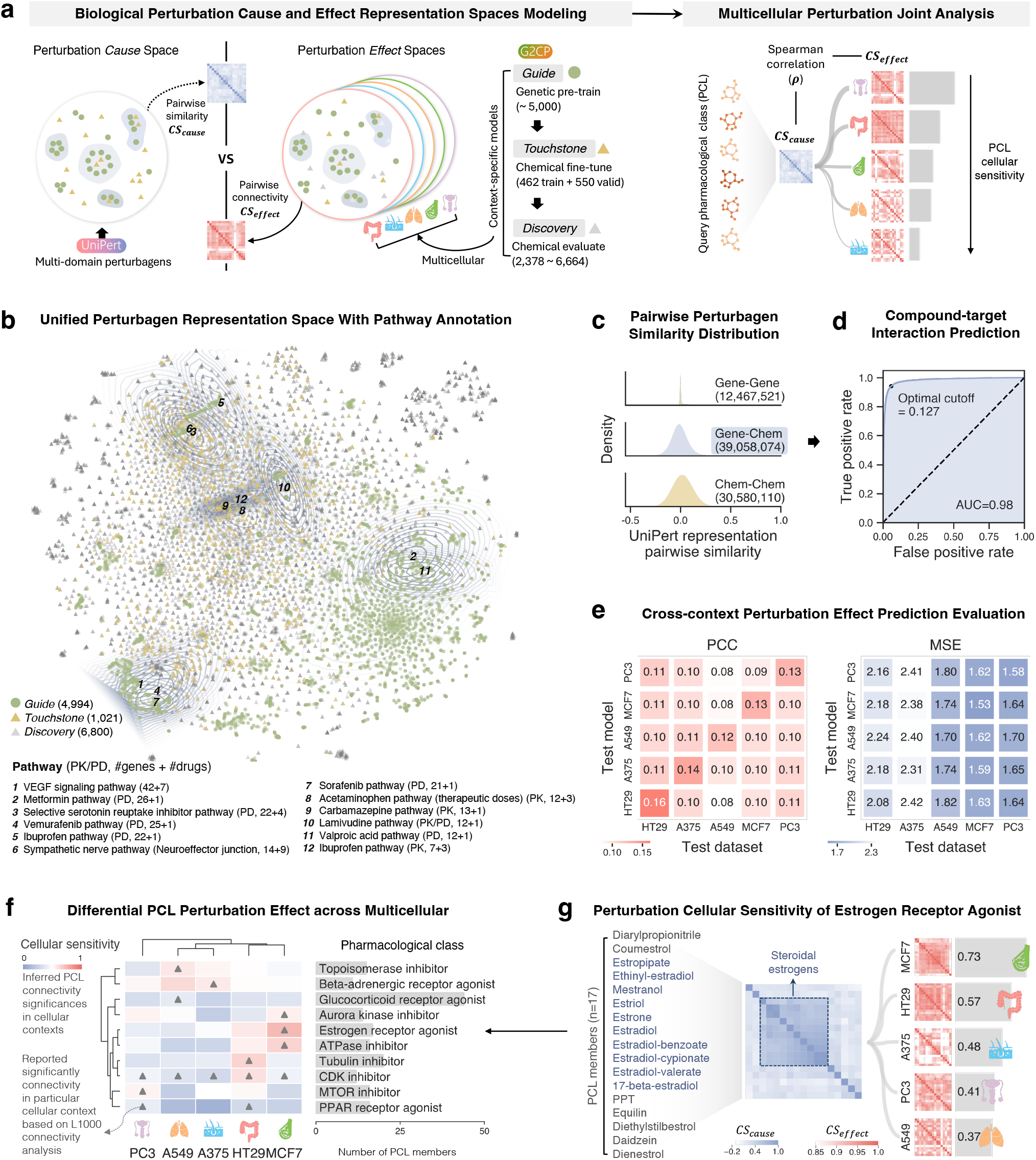
UniPert-G2CP reveals PCL cellular sensitivity through generated causal representations. **(a)** Workflow for characterizing pharmacological class (PCL) cellular sensitivity using UniPert-G2CP. Given defined *Guide, Touchstone, Discovery* perturbagen sets, UniPert generate perturbation cause embeddings, while G2CP trained multiple cancer cell line models in a cost-efficient way and simulate cellular phenotypic effects of unseen perturbation, resulting multiple causal representation spaces (left). For a PCL with multiple compound members, stronger correlations between their pairwise cause similarity (*CS*_*cause*_) and context-specific effect connectivity (*CS*_*effect*_) matrix indicates more consistent responses induced in the particular context, resulting a higher cellular sensitivity (right). **(b)** t-SNE plot visualizing UniPert embeddings for perturbagens from the *Guide* (4,994 genes, green circles), *Touchstone* (1,021 compounds, yellow triangles), and *Discovery* (6,800 compounds, gray triangles) sets. Overlaid contour density plots represent pathway annotations collected from the PharmGKB resource (PK, pharmacokinetics; PD, pharmacodynamics). **(c)** Distribution of pairwise UniPert embedding similarities across approximately 82 million perturbagen pairs. Similarities are grouped into gene-gene, gene-chem, and chem-chem categories, derived from 4,994 genetic (*Guide*) and 7,821 chemical perturbagens (*Touchstone* and *Discovery* combined). **(d)** ROC curve assessing the ability of gene-chem pairwise cosine similarity scores from UniPert embeddings to recover known interactions. Performance is evaluated against 81,397 experimentally validated compound-target interactions used as positive references. **(e)** Cross-cell-line evaluation on five cell line-specific models trained using C2CP. These perturbation effect prediction models were trained under the same perturbation conditions. Each cell line test set consists of a different number of compounds. **(f)** Characterized differential PCL cellular sensitivities. Heatmap shows inferred cellular sensitivity scores for multiple pharmacological classes across the five cell lines. Triangle markers indicate significant connectivity reported from L1000 analysis. Bar chart (right) displays the number of members per PCL. **(g)** Cellular sensitivities analysis for the estrogen receptor agonist PCL across 5 cancer cell lines.

We first constructed a context-free, cross-domain perturbagen embedding space using UniPert for the *Guide, Touchstone*, and *Discovery* sets (Supplementary Table 4). By integrating PharmGKB pathway evidence^88^, including manually curated relationships between drugs and genes involved in pharmacokinetics (PK) and pharmacodynamics (PD), we annotated this space. T-SNE visualizations and contour plots based on pathway labels reveal that UniPert effectively clusters genes and drugs involved in shared pathways (Figure 5b). Notably, certain pathways exhibited similar patterns, such as the selective serotonin reuptake inhibitor pathway and the sympathetic nerve pathway, as well as the Vemurafenib and Sorafenib pathways. UniPert also distinguished between the PK and PD roles of the same compound, exemplified by Ibuprofen, which formed separate contours according to its mechanistic context.

To quantify perturbagen relationships within this unified embedding space, we computed pairwise cosine similarity scores across all pairs from 4.994 genetic perturbagens (gene) and 7,821 chemical perturbagens (chem), resulting in a perturbagen association matrix. Among over 82 million pairs, gene-gene and chem-chem pairs displayed markedly different distributions (Figure 5c). Chem-chem similarities followed a broader, near-normal distribution, reflecting the structural diversity and flexibility of small molecules. In contrast, the gene-gene similarities were more narrowly distributed, which can be attributed to the evolutionary conservation and functional constraints of biomolecules. The similarity distribution of gene-chem pairs closely mirrored that of the chem-chem pairs but with a slightly lower mean, suggesting that UniPert coherently represents diverse perturbagen types into a unified semantic space.

To further validate whether the UniPert-based perturbagen similarity matrix can recover known compound-target relationships, we compared approximately 40 million gene-chem perturbagen pairs against experimentally supported 81,397 compound-target interactions used in UniPert training. Labeling these interacting pairs as positive samples and the rest as negative samples, we performed receiver operating characteristic (ROC) analysis. The resulting area under the curve (AUC) exceeded 0.98, with an optimal cutoff of 0.127, confirming the efficiency and promise of UniPert embeddings for similarity-based compound– target prediction (Figure 5d). In previous studies of cross-domain perturbation connection analyses^5,81,89,90^, perturbagen associations were manually curated through literature-based annotations of shared descriptors, such as MOA, GO, and pathway membership. However, this process i) is cumbersome and time-consuming, ii) produces binary values that fail to reflect the strength of relationships, and iii) heavily relies on prior knowledge, limiting its applicability to understudied targets and unannotated compounds. UniPert’s multimodal, sequence-based molecular representation method overcomes these limitations effectively.

Last, we performed cross-context validation on *Discovery* sets using five trained predictors to evaluate whether they can capture the heterogeneity of perturbation responses across diverse cellular contexts, which mimics the biological complexity encountered in precision medicine applications (Figure 5e). The resulting PCCs demonstrated that each cellular model exhibited the expected strongest performance on its corresponding test set, highlighting the ability of these perturbation phenotype predictors to capture underlying cellular heterogeneity. While for MSE, although each model also performed optimally on its matched context, the differences across datasets were more pronounced. These findings reinforce our earlier observation that the direction of perturbation-induced transcriptional changes is more robust and predictable than their magnitude.

### *Cause-Effect* Space Joint Analysis Characterizes PCL Perturbation Cellular Sensitivity

Mechanism-driven therapy offers a critical avenue for advancing personalized medicine, particularly by leveraging PCL-level knowledge to guide therapeutic decisions^91^. A drug typically belongs to a single PCL but may span multiple therapeutic indications. For instance, metoprolol is clinically used as both an antianginal and an antihypertensive agent through β-adrenergic receptor blockade^92^. Although compounds within a PCL share a common MOA, their induced phenotypic responses are often modulated by the cellular context in which they act. Such systematic differences in transcriptional response across disease-relevant cellular environments can translate into distinct therapeutic outcomes. Therefore, quantifying the cellular response sensitivity of a defined PCL is essential for uncovering context-aware therapeutic vulnerabilities and informing precision treatment strategies^93^.

We compared UniPert-G2CP-derived perturbation cause and effect spaces to measure this cellular sensitivity. For each PCL, we computed two types of distance matrices: i) a context-independent molecular similarity matrix (CS_*cause*_) based on UniPert embeddings, and ii) a set of context-dependent phenotypic connectivity matrices (CS_*effect*_) predicted by previously trained cellular perturbation simulators across five cancer cell lines (Figure 5a, left). CS_*cause*_ provides a generalizable baseline for comparing phenotypic connectivity across diverse contexts, thereby obviating the need for multi-context normalization. We employed Mantel tests with Spearman rank correlation coefficient (*ρ*, ranging from -1 to 1) to quantify the correlation between CS_*cause*_ and CS_*effect*_ (Figure 5a, right). Higher correlations indicate stronger cellular sensitivity, suggesting that the transcriptional responses to PCL members are more robust and consistent within particular cellular contexts.

We conducted this joint analysis on a curated set of experimentally validated PCLs with sufficient membership from *Subramanian et al*^5^. As expected, most PCLs exhibited strong inter-member molecular structural similarity and context-dependent phenotypic connectivity (Extended Data Figure 8). All PCLs yielded measurable (*ρ* > 0) while varying perturbation effect sensitivities across five contexts (Figure 5f). Notably, estrogen receptor (ER) agonists, a key mediator of estrogen signaling in tissues such as the breast, exhibited pronounced context dependency, with a most significant sensitivity observed in MCF7 breast cancer cells (*ρ* = 0.73; Figure 5g). Similarly, other examples such as the peroxisome proliferator-activated receptor (PPAR) agonists showed maximal sensitivity in HT29 and PC3, which is also consistent with prior reports. Furthermore, we also observed that CS_*cause*_ of ER agonists captured a remarkable structural similarity pattern among steroidal estrogens that share a conserved tetracyclic steroid scaffold (Figure 5g). Among them, 17-beta-estradiol exhibited the highest structural similarity to other members and is also known as one of the most potent ER agonists. However, this steroidal estrogen pattern was less pronounced in the CS_*effect*_ matrices, confirming that phenotypic convergence should not only be driven by chemical structure similarity but strongly influenced by cellular environment.

UniPert-G2CP enable systematic quantification of context-specific therapeutic signatures from constructed perturbation *cause-effect* representation spaces. This capacity offers latent therapeutic opportunities that might be overlooked by structure- or phenotype-based analyses alone. Ultimately, our joint modeling and analysis of biological perturbations across multimodal, multi-domain, and multicellular contexts establishes a new AI-driven paradigm for precision pharmacology strategies that are both mechanism-informed and context-aware, providing a promising path toward more rational and individualized therapeutic interventions.

## Discussion

Despite considerable progress in modeling perturbation-induced cellular states^94–96,66,67,2^, most existing methods have focused primarily on phenotypic effect prediction while neglecting the semantic structure and relations inherent to perturbation agents. Moreover, significant differences in data modality and scale between perturbations from cell-intrinsic and -extrinsic factors have impeded the cross-domain modeling and integrative analysis. UniPert-G2CP addresses these limitations by unifying perturbation representations from both cause and effect perspectives^97,98^, leveraging multimodal molecular representation learning and cross-domain transfer learning. This approach substantially enhances the generalizability of biological causal models across data modalities, experimental scales, biological contexts.

At the molecular (perturbation cause) level, UniPert establishes semantic relations between large biomolecules and small molecules into a shared embedding space using prior knowledge and contrastive learning. The encoders integrate ECFP, MSA, PLM, and GNN to preserve meaningful functional group information across multimodal molecules. This interpretable, scalable space enables pharmacological functional annotation and clustering of understudied proteins and drug-like compounds. Serving as a biological knowledge engine, this molecular semantic bridging not only improves the prediction of transcriptional responses to previously unseen gene perturbations or compound treatments, but also alleviates the need for manual annotation, facilitating cross-domain screen analyses, streamlines computational workflows, and expands the scope of perturbation study.

At the phenotypic (perturbation effect) level, the G2CP framework transforms the cost-efficiency of *in silico* high-content drug screening by utilizing dense, systematic genetic screens to compensate sparse chemical screens with vast search spaces. Extensive experiments demonstrate that the “genetic pre-training + chemical fine-tuning” framework efficiently predicts compound responses in multiple cellular contexts, capturing context-dependent drug effects that would otherwise require extensive experimental manipulation to observe. Although current screening efforts primarily focus on transcriptomic profiling, image-based phenotypic screens hold considerable potential for future model development as its more accessible, less expensive, higher content and throughput cellular profiling^90,99–101^. Moreover, as screening data from additional omics layers^102,103^, e.g., such as proteomics^104–106^ and epigenomics^107–109^, continue to grow, G2CP is well positioned to support integrative modeling of multi-omics cell states.

From the standpoint of computer-aided drug discovery, UniPert-G2CP advances a shift in drug virtual screening from traditional cellular context-free approaches such as docking-based methods^110,111^, toward a unified framework that jointly models context-independent molecular relationships alongside context-dependent phenotypic consequences^112^. This biologically grounded strategy facilitates more realistic *in silico* compound prioritization and early-stage screening across vast chemical spaces, bridging molecular-level modeling with preclinical drug response prediction.

In the context of translational precision medicine, UniPert-G2CP paves the way for *in silico* clinical trial simulations and personalized treatment planning by predicting individual drug responses based on patient-specific genetic backgrounds^3,113,114^. Moreover, by disentangling perturbation effects across diverse cellular contexts, UniPert-G2CP provides a powerful tool for systematically investigating mechanism-aware drug response heterogeneity. These phenotypic analogies provide a foundation for identifying new therapeutic indications and support mechanism-informed drug repurposing strategies beyond conventional target-centric approaches^115^.

While UniPert-G2CP unifies genetic and chemical perturbations, it cannot explicitly represent different layers of genetic perturbations, as UniPert relies on the protein-layer sequence to encode genetic interventions. Consequently, it cannot represent perturbations targeting non-coding regions that play critical roles in gene regulation^116,117^. Future extensions could incorporate biosequence language models across multiple layers—such as Evo^29^ for DNA and RNAErnie^118^ for RNA—to provide more fine-grained and comprehensive representations for genetic perturbations.

At its core, UniPert-G2CP is the first universal framework biological perturbation modeling across genetic and chemical domains, marking a computationally validated conceptual advance toward biological causal foundation model construction and AI virtual cells biulding^1,2,35,36,119^. It lays the groundwork for robust, data-driven prediction frameworks that can inform personalized treatment strategies and accelerate the realization of precision medicine. We anticipate that UniPert-G2CP will make a significant contribution to the field of perturbation biology and advance the frontiers of drug discovery and translational therapeutics.

## Methods

The Methods describe in turn i) the model architecture, training strategy, and implementation details of UniPert and G2CP, ii) the implementation for genetic and chemical perturbation effect prediction tasks, iii) all datasets used for model training and downstream applications, and iv) all statistical analyses and evaluation methods used.

### UniPert model

#### Model architecture

A schematic overview of UniPert is depicted in Extended Data Figure 1b. UniPert integrates chemical and genetic perturbagen encoder modules into a unified, sequence-based, end-to-end representation learning framework.

#### Chemical perturbagen encoder module

UniPert first characterizes genetic perturbagens using the topology-based ECFP algorithm^30^. Specifically, for a given small molecule **c**, UniPert employs the RDKit tool^120^ to calculate the molecular fingerprint, generating an initial *K*-dimensional binary vector **h**_**c**_ ≔ ℱ_*ECEP*_(**c)** ∈ {0,1}^*K*^. The vector is then passed through a fully connected layer and transformed into a *D*-dimensional dense vector: 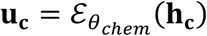,where *θ*_*chem*_ represents the learnable parameters of the chemical perturbagen encoder 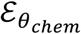,and **u**_**c**_ ∈ ℝ^*D*^ denotes the generated UniPert embedding for the query **c**.

#### Genetic perturbagen encoder module

UniPert encodes genetic perturbagens through a combination of sequence alignment algorithm, pre-trained PLM, and GNN. Specifically, UniPert first constructs a reference graph **G** = **(V, E)**, where **V** denotes the protein node set of the human genome-wide proteins and their corresponding sequences (*N* = 19,187) downloaded from the HGNC^121^ and UniProt^122^ resources, and **E** is the weighted edge set representing pairwise Smith-Waterman-based^123^ sequence alignment similarity scores between proteins, calculated using MMseqs2^124^. Given an unseen amino acid sequence **t**, UniPert adds it as an new node to **G**, resulting in a update graph **G**_**t**_ ≔ **(V**_**t**_, **E**_**t**_), where **V**_**t**_ = **V** ∪ {**t**} and **E**_**t**_ = **E** ∪ **e**_**t**_, with **e**_**t**_ representing the new edges between **t** and the nodes in **V** based on their sequence similarities. To obtain and update the embedding for **t**, UniPert employs the ESM model (esm2_t33_650M_UR50D version)^27^ to initialize the node embeddings for **V**_**t**_ as *P* -dimensional (*P* = 1280) vectors, resulting in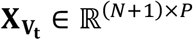. Subsequently, a two-layer GNN is applied to perform message passing on **G**_**t**_, updating the node embeddings to *D*-dimensional vectors, 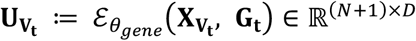,where *θ*_*gene*_ represents the learnable parameters of the genetic perturbagen 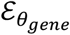,and 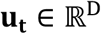 extracted from 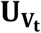 represents the generated UniPert embedding for the query **t**.

### Training strategy

UniPert employs a dual optimization strategy to effectively train the model, including graph self-supervised learning for genetical perturbagen embedding enhancement and contrastive learning for genetical-chemical perturbagen alignment, performing the overall loss *ℒ* = *ℒ*_*enhance*_ + ℒ_*align*_

### Graph self-supervised learning los

Inspired by the successful applications of bootstrap mechanisms in representation learning within the fields of computational vision^125^ and large-scale graphs^54^, UniPert adopts a bootstrapped graph latents approach^54^ for efficient self-supervised representation learning of genetic perturbagens. Specifically, for the constructed reference graph **G** = (**X, A)** with initialized ESM node features **X** ∈ ℝ^N×P^ and similarity-based adjacency matrix **A** ∈ ℝ^*N×N*^, UniPert produces two augmented graph variants: 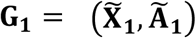 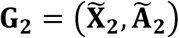,by applying stochastic masking to edges and node features. These variants are processed separately through an online encoder 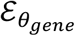 and an offline encoder 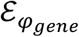,producing corresponding node representations 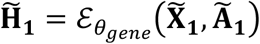 and 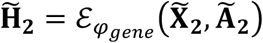.The online representations are then fed into a node-level predictor 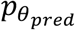,yielding 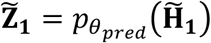 aimed at predicting the offline representations 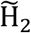,with a cosine similarity-based loss *ℒ*_enhance_(*θ, φ*) defined as

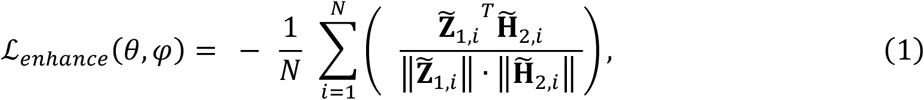

where N is the number of graph nodes, and 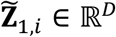 and 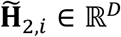 represent the embeddings of the *i*-th node from the online predictor 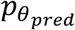 and offline encoder 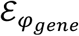,respectively. During training, the online parameters *θ* are updated through gradient descent, while the offline encoder parameters *φ* are stop-gradient and updated as an exponential moving average of *θ*_*gene*_ at each step. This approach alleviates the need for numerous negative samples and ensures stable and efficient learning on the large-scale protein graph.

### Contrastive learning loss

To establish functional and mechanistic relationships between genetic and chemical perturbagens, UniPert adopts a contrastive learning strategy^126^ to align representations of paired perturbagens (e.g., a compound and its protein target). Specifically, given a minibatch of *M* experimentally validated compound-target interactions, the representations of the *i*-th pair (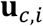 and 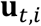) are encoded as 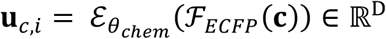. and 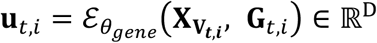,respectively. To promotes a dense clustering of positive pairs in the embedding space, UniPert optimizes the symmetric contrastive loss *ℒ*_*align*_

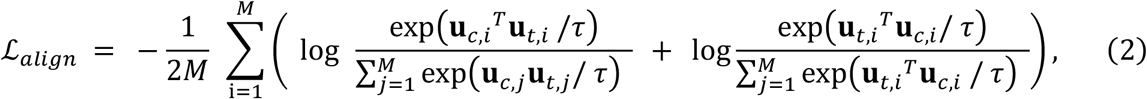

where *τ* > 0 is the temperature parameter that regulates the similarity distribution. This alignment ensures embedding coherence across molecules at different scales, enhancing the biological interpretability and robustness of the unified representation space.

### Implementation details

We implemented UniPert training using PyTorch (Version 1.12.1)^127^ and PyTorch Geometric (Version 2.5.2)^128^ on a single NVIDIA Tesla V100-SXM2-32GB GPU. Optimal hyperparameters were determined through grid search for UniPert training, with selection based on the comprehensive performance ranking across *ℒ*_*enhance*_ and *ℒ*_*align*_ losses. TensorBoard^129^ was employed for real-time tracking and visualization of training performance.

To facilitate direct inference of representations from new query gene or compound names using our trained UniPert model, we also integrated the getSequence^130^ and PubChemPy^131^ packages to encapsulate UniPert. This integration allows UniPert to retrieve corresponding amino acid sequences or SMILES strings via online API interfaces and then generate perturbagen embeddings.

### G2CP framework

#### Framework architecture

The cross-domain perturbation transfer learning ability of G2CP primarily benefits from UniPert’s prior knowledge-enhanced and multimodal perturbagen encoders. Thus, G2CP can be seamlessly integrated into any perturbation effect prediction model architecture that takes perturbation conditions and phenotypic profiles as input. In this study, we used GEARS^66^ as the backbone model to implement G2CP on the LINCS and CPJUMP1 datasets.

#### Training strategy

A two-stage *genetic-to-chemical* perturbation transfer learning strategy (illustrated in Figure 4b) is designed to train a contextual perturbation effect prediction model 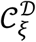 with learnable parameters *ξ* for accelerating large-scale *in silico* drug screening by effectively transferring response from perturbagen genetic to chemical perturbagens with UniPert encoders 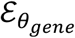 and 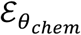.Specifically, given a context-specific dual-domain perturbation dataset 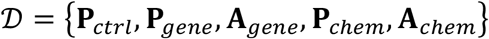,containing *L* –dimensional unperturbed (control group) phenotype 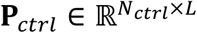,post-genetic-perturbation phenotype profiles 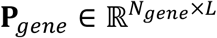,post-chemical-perturbation 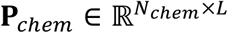,along with their corresponding genetic and chemical perturbagen annotations 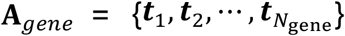 and 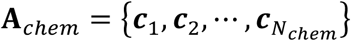.

#### Pre-training stage

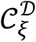 is initially trained using genetic perturbation data, 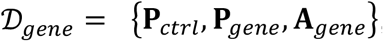,which, due to advancements in current screening technologies, usually owns a more variety of perturbagen types compared to lab-produced chemical perturbation data. The model predicts post-genetic-perturbation phenotype profiles as 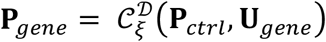, where 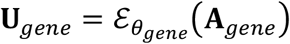 represent generated genetic perturbagen embeddings by UniPert. Following pre-training, the model 𝒞_*ξ*_ systematically learns the genetic characteristics of the cellular environment and the response patterns to perturbagens targeting distinct molecular pathways. This enables it to effectively predict phenotypic effects of genetic perturbations and establish a robust foundation for subsequent chemical perturbation simulation.

#### Fine-tuning stage

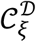 continues to be trained using chemical perturbation data 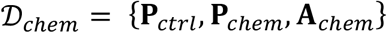, following 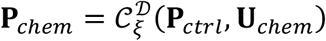,where the chemical perturbagen embeddings 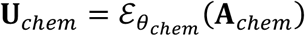 are generated with UniPert. Leveraging the pre-trained knowledge from genetic perturbations, 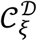 is further fully fine-tuned to accurately model chemical perturbation responses, thereby effectively transferring the learned perturbation dynamics across domains to enhance prediction performance.

### Implementation for genetic perturbation effect prediction task

To evaluate the performance of different genetic perturbagen encoders on the task, we employed the GEARS-derived framework (illustrated in Extended Data Figure 5a). We compared and evaluated the impact of various encoders, including the original GEARS^66^ encoder, PseAAC^68^, ESM^27^, OntoProtein^49^ and UniPert. Apart from the original GEARS re-learns perturbagen representations for each dataset training based on prior GO knowledge, other methods rely on pre-trained/pre-defined models that capable of directly obtaining sequence-derived features and generalize to any amino acid sequence. These encoding methods are described below:

### PseAAC^68^

A widely used pseudo amino acid composition method that considers the sequential effects of amino acids in a protein. We utilized the propy3 package^132^ to calculate 68-dimensional PseAAC embeddings with the parameter *λ* set to 24 to accommodate the shortest sequence length of 24 in the reference protein library.

### ESM^27^

A transformer-based large-scale pre-trained protein language model that generates 1280-dimensional informative embedding (esm2_t33_650M_UR50D version) with evolutionary patterns for each given sequence.

### OntoProtein^28^

A knowledge-enhanced protein pre-training model incorporating GO knowledge graph information that generate 1024-dimensional embeddings for each sequence.

These methods generate embeddings of varying dimensionalities, which are subsequently mapped to a unified dimension through fully connected layers for a fair comparison. The training hyperparameter settings used for genetic perturbation effect prediction tasks are consistent with the original GEARS study^66^.

### Implementation for chemical perturbation effect prediction task

To benchmark the impact of different compound encoders on the prediction of phenotypic outcomes following drug treatments, we employed a CPA-derived framework (illustrated in Extended Data Figure 6a). The comparative encoders including ECFP^30^ (used in chemCPA^67^), Uni-Mol^31^, KPGT^32^, and UniPert,, all of which are capable of representing unseen compounds. The details of each encoder are described as follows:

### ChemCPA^67^

Following the tutorial of extended CPA to predict unseen compound perturbation response, we utilized the RDKit tool^39^ to compute 2048-dimensional ECFP features with a radius of 2 for each given small molecule.

### Uni-Mol^31^

A transformer-based 3D molecular representation model that produces 512-dimensional embedding for each small molecule.

### KPGT^32^

A knowledge-guided pre-trained graph transformer model that incorporates molecular graph structures and external knowledge to generate 2304-dimensional embeddings for each small molecule.

The training hyperparameter settings for CPA-derived chemical perturbation effect prediction training with all encoders followed the CPA tutorial^67^.

### Datasets

#### Compound-target interaction dataset

We curated a comprehensive compound-target interaction dataset according to the involved compounds of the CMAP project^5^. This dataset was gathered from the PubChem resource^133^ and consists of 81,397 pairs across 8,533 compounds and 4,250 proteins, integrated from 8 related compound-target interaction databases^134–141^. The corresponding SMILES sequences and amino acid sequences were collected for UniPert model training^122^.

#### Drug and protein target annotations

Classifications of small-molecule drugs and protein targets were collected from the ChEMBL database^141^, including 2,812 drugs across 928 MOA classes, and 4,417 protein targets annotated with multi-level hierarchical pharmacological classes (PCLs). The top 18 MOA categories used for drug embedding semantic similarity analysis cover 726 drugs. Detailed information on full class names, abbreviations and class hierarchy is provided in Supplementary Tables 1 and 2. Protein subfamily/family/superfamily classifications were collected from the UniProt resources^122^.

#### *Dixit et al*. dataset

The *Dixit et al*. dataset is a single-gene perturbation dataset downloaded from the processed data provided by the GEARS study^17,66^, including 44,375 profiles about 20 genes perturbing the K562 cell line.

#### *Norman et al*. dataset

The *Norman et al*. dataset is a combination genetic perturbation dataset, also downloaded from the processed data available through the GEARS study^142,66^, comprising 91,205 profiles from 283 single and combination genes perturbing the K562 cell line.

#### Sci-Plex3 dataset

The sci-Plex3 dataset is a large-scale chemical perturbation dataset involving 187 drugs and 3 cell lines, downloaded from the processed data from the CPA study^24,67^. It was used to evaluate the performance in predicting perturbation effects for drugs held out in pathway-based hierarchical splits.

#### HDAC inhibitor dataset

The HDAC inhibitor dataset is a small-scale chemical perturbation dataset extracted from the sci-Plex3 dataset for leave-one-drug-out evaluation. It includes phenotype data from both the untreated control group and cells treated with 17 HDAC inhibitors at 1 µM for 24 hours across 3 cell lines.

#### CPJUMP1 datasets

The CPJUMP1 datasets^81^ include dual-domain perturbation data for two cell lines, i.e., A549 and U2OS. We downloaded pre-processed well-level morphological profiles, with each profile containing 838-dimensional features generated by the image feature extraction software CellProfiler^143^ and the pre-processing tool Pycytominer^144^, resulting the CPJUMP1-A549 and CPJUMP1-U2OS datasets with detailed information in Supplementary Table 3.

#### LINCS datasets

We extracted transcriptome-based perturbation data from the CMap LINCS resource^5^ (level 5 signatures of LINCS2020 Release) for cell lines that underwent both CRISPR knockout and compound screening. These data were subjected to several quality control and pre-processing steps, including batch correction, differential expression normalization, and replicate consensus processing, which ensured greater robustness and more significant perturbation effects compared to the original readouts. We selected cell lines with the top 5 number of perturbagens under specific conditions (CRISPR sgRNA infection for 96h and compound treatment at 10uM for 24h), forming five cell-line-specific dual-domain perturbation datasets, i.e., LINCS-A375, LINCS-A549, LINCS-HT29, LINCS-MCF7, LINCS-PC3, with detailed information in Supplementary Table 3.

### Evaluation methods and metrics

#### Intra-MOA drug embedding semantic similarity analysis

To quantify and compare the compound (chemical perturbagen) semantic distance within MOA classes based on ECFP fingerprints and UniPert-generated embeddings, we first computed both Tanimoto coefficient and cosine similarity for 3,952,266 pairs of all collected drugs (*n*_*all*_ = 2,812), respectively. The Tanimoto coefficient is widely regarded as the most appropriate metric for fingerprint-based similarity calculations^145^, and defined as

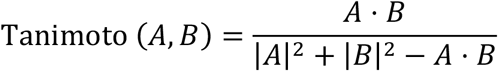

where *A* and *B* represent the binary fingerprint vectors of two compounds. The cosine similarity is commonly used to assess vector-based embedding similarity in deep learning models and computed as

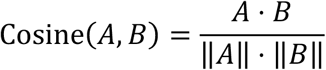

where *A* and *B* denote the continuous-valued embedding vectors of the compounds, and ‖*A*‖ is the Euclidean norm of vector *A*.

Since the numerical ranges of similarity measures differ—typically [0, 1] for Tanimoto and [– 1, 1] for cosine—we employed the standardized mean difference (SMD) to quantify the similarity divergence between each MOA group (comprising intra-MOA drug pairwise similarities) and the overall group (consisting of all pairwise similarities), as follows:

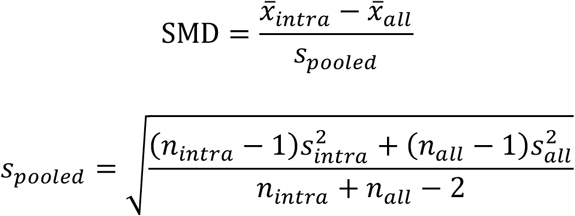

where 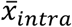 and 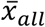 denote the mean similarity values within the given MOA group and the overall group, respectively; 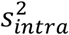 and 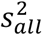 are their corresponding standard deviations; and 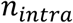 represent the number of pairwise comparisons within the MOA group. By normalizing the mean difference against the pooled standard deviation, SMD enables the direct comparison of similarity distributions derived from heterogeneous metrics. A larger SMD reflects stronger intra-class consistency of similarity relative to the overall distribution, suggesting a greater ability of the encoder to capture the semantic similarity among drug members within the MOA class.

#### Protein Target embedding pharmacological clustering analysis

To evaluate the pharmacological interpretability of protein target embeddings generated by UniPert, we conducted clustering analysis on 4,417 human protein targets annotated with pharmacological labels organized across multiple levels of class granularity. Specifically, we analyzed 35 classed across 5 levels of annotation hierarchy, with the hierarchical relationships illustrated in Extended Data Figure 2a). For each parent class, we first extracted the UniPert embeddings of all protein target members and performed K-means clustering to evaluate their semantic coherence within the embedding space. Each K-means clustering was applied with the number of clusters corresponding to the unique labels at each annotation level.

The clustering performance was quantitatively assessed using two standard metrics: the Adjusted Rand Index (ARI) and the Normalized Mutual Information (NMI), which measure the concordance between predicted cluster assignments and ground-truth class labels. The ARI measures pairwise agreement between predicted and true labels while adjusting for chance, and is defined as:

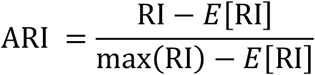

where RI denotes the Rand Index and *E*[RI] is its expected value under random labeling. The NMI quantifies the mutual dependence between the predicted clusters *C* and the ground-truth class labels *L*, normalized by the geometric mean of their individual entropies:

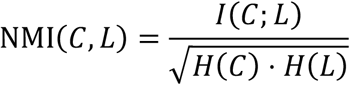

where *I*(*C*; *L*) is the mutual information between *C* and *L*, and *H*(*C*) and *H*(*L*) denote the Shannon entropy of the predicted clusters and true labels, respectively. Higher scores in both metrics indicate stronger alignment between known pharmacological classifications and embedding-inferred clusters across different target encoding methods.

To visualize class separability in the high-dimensional embedding space, we used t-distributed stochastic neighbor embedding (t-SNE) to project the embeddings into two dimensions and colored the points according to their subclass labels at each parent class.

#### Protein functional conserved motif enrichment analysis

To assess the fine-grained representational capacity of different protein sequence encoders in capturing biologically meaningful local sequence patterns, we performed motif enrichment analysis using the AME tool^146^ in the MEME Suite (v5.5.7)^147^ on the reference set of 19,187 human proteins. We conducted motif enrichment testing against known protein sequence motifs curated from two publicly available databases: the Eukaryotic Linear Motif (ELM) and PROSITE databases. The reference sequences were scanned for overrepresented conserved motifs. Fisher’s exact test was used to assess motif enrichment, statistical significance was evaluated by E-values, and the sequence scoring method was performed using the average odds score. Top significantly enriched motifs were selected, and the UniPert, ESM, and OntoProtein embeddings of proteins containing each motif were extracted for t-SNE visualization. Proteins associated with the same motif were highlighted to assess whether different encoders could group proteins sharing conserved functional subsequences into coherent clusters within the embedding space.

#### Perturbation effect prediction evaluation

In the genetic and chemical perturbation effect prediction tasks for OOD conditions, we used two major complementary metrics to evaluate prediction performance: Pearson correlation coefficient (PCC) and mean squared error (MSE). PCC captures the consistency of perturbation response patterns by measuring the linear correlation between observed and predicted profiles, whereas MSE emphasizes the accuracy of reproducing measured effects by quantifying absolute differences.

For each test perturbation condition *c*, we compared the mean observed post-perturbation profile 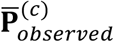 and the mean predicted post-perturbation profile 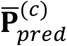,where each profile is represented as a *d* -dimensional vector corresponding to the expression values of *d* genes. The PCC and MSE 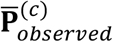 and 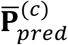 are defined as:

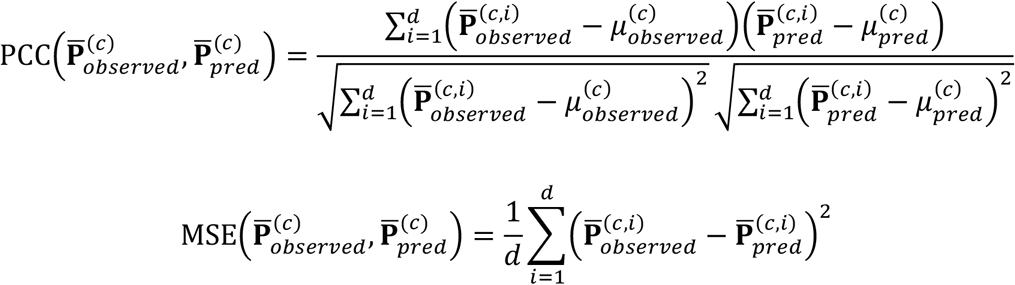

where 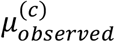 and 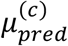 denote the means of 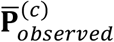 and 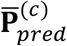,respectively. Conceptually, PCC reflects whether the predicted directions of gene expression changes align well with observations, while MSE focus more on the precision of the predicted magnitudes. By combining the two metrics, we can assess model’s predictive ability from different perspectives. Additionally, the *R*^2^ metric is also used for evaluation of perturbation predictions, computed as:

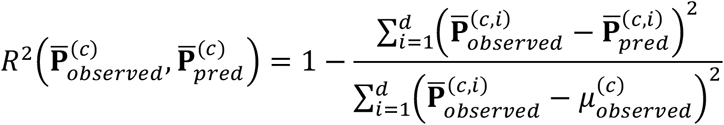

The overall evaluation for each OOD test set is the average of the evaluations across all test conditions.

#### Multi-domain perturbagen (perturbation cause) similarity calculation

We used cosine similarity to measure the distance between molecular pairs of gene-gene (n = 12,467,512), chem-chem (n = 30,580,110), gene-chem (n = 39,058,074) of perturbagens in the *Guide* (n = 4,994), *Touchstone* (n = 1,021), and *Discovery* (n = 6,800) sets within the UniPert-derived shared embedding space.

#### Embedding similarity-based compound-target interaction prediction evaluation

To assess the compound-target interaction prediction capability based on the UniPert embeddings, we performed a receiver operating characteristic (ROC) analysis on gene-chem 39,058,074 pairs, involving 4,994 proteins and 7,821 compounds. We spitted all possible pairs into two collections serving as their true labels: expected positive pairs (*E*_*p*_), which were known to interact in our previous collected compound-target interaction dataset (n = 81,397), and null negative pairs (*N*_*p*_), which were not recorded as interacting. The prediction score for each compound-target pair was derived directly from the calculated UniPert embedding-based gene-chem similarity score, where higher similarity was indicative of a higher likelihood of interaction. By varying the discrimination threshold applied to these similarity scores, we generated the ROC curve, plotting the true positive rate (TPR) against the false positive rate (FPR) at various threshold settings. The performance of the prediction method was quantified by calculating the area under the ROC curve (AUC). The optimal discrimination threshold was determined by maximizing the Youden’s J statistic (TPR−FPR).

#### Cellular phenotype (perturbation effect) connectivity calculation

To quantify the phenotypic similarity between two post-perturbation profiles **P**_*i*_ and **P**_*j*_ of perturbagens *i* and *j*, we calculated their connectivity based on the weighted connectivity score employed in the CMap analyses^5^, which is a composite, bi-directional version of the weighted Kolmogorov-Smirnov enrichment statistic (ES)^148^. The connectivity between **P**_*i*_ and **P**_*j*_ is calculated as follows:

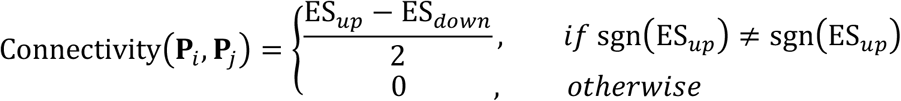

where *ES*_*up*_ is the enrichment of 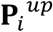 in **P**_*j*_ and *ES*_*down*_ is the enrichment of 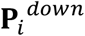 in **P**_*j*_. Genes with positive scores are considered upregulated and those with negative scores are considered downregulated in our inference perturbated profiles of the LINCS datasets. This approach allows us to assess the similarity in perturbation-induced phenotypic responses across various perturbagen conditions, effectively capturing the connectivity of perturbagen effects in a given cellular context.

#### PCL cellular sensitivity calculation

The joint analysis and comparison of the perturbation *cause-effect* spaces, comprising UniPert-generated context-free molecular perturbagen embeddings and G2CP-predicted context-aware phenotypic state representations, enables systematic investigation of cellular response significance across diverse contexts within PCLs. To quantify the cellular sensitivity of a given PCL *p*, we first computed its pairwise drug similarity matrix 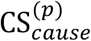 based on UniPert embeddings. 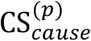 provides a baseline for understanding how drugs within the same PCL are related in a molecular sense, independent of cellular context. For each specific cellular context *c*, we then computed the corresponding pairwise connectivity matrix 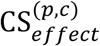 based on G2CP-predicted perturbation profiles. These context-aware matrices characterize the degree of phenotypic coherence among drugs within the same MOA under specific cellular contexts.

We subsequently hypothesized that the cellular context in which the context-aware phenotypic connectivity matrix 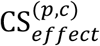 most closely aligns with the molecular similarity matrix 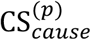 reflects the strongest and most coherent perturbation responses. This hypothesis is grounded in the findings of *Subramanian et al*.^5^, which demonstrated that perturbation phenotypic connectivity can be indicative of context sensitivity. To quantify this alignment, we employed the Mantel test, a statistical method commonly used to assess the correlation between two distance or similarity matrices. The Mantel test yielded a Spearman correlation coefficient *ρ*^(*p,c*)^ and a nominal two-sided p value. A higher *ρ*^(*p,c*)^value indicates a higher degree of cellular sensitivity for PCL *p* in the context *c*. The p value was estimated via 1,000 permutations and reflects the statistical significance of the observed correlation against the null hypothesis of no association between the matrices. Comparing *ρ*^(*p,c*)^across cellular contexts thus enabled the identification of those in which a PCL’s molecular coherence most strongly translates into consistent phenotypic responses.

## Data availability

All datasets used in this work were obtained from public repositories. The human genome-wide proteins, as well as their corresponding sequences and family annotations were obtained from https://www.genenames.org/ and https://www.uniprot.org/. The compound-target interaction dataset was obtained from https://pubchem.ncbi.nlm.nih.gov/. The annotations of hierarchical target classes and drug MOA classes were obtained from https://www.ebi.ac.uk/chembl/. The *Dixit et al*. and *Norman et al*. datasets were obtained from https://github.com/snap-stanford/GEARS. The sci-Plex3 dataset was obtained from https://github.com/theislab/cpa. The CPJUMP1 datasets were obtained from https://github.com/jump-cellpainting. The LINCS datasets were obtained from https://clue.io/data/CMap2020#LINCS2020. The pathway annotations were obtained from https://www.pharmgkb.org/. The pharmacologic class and membership, as well as the reported PCL context sensitivity information was obtained from https://www.cell.com/cell/fulltext/S0092-8674(17)31309-0.

## Code availability

UniPert is available at https://github.com/TencentAILabHealthcare/UniPert, which provides user-friendly tutorials for encoding genetic and chemical perturbagen condition information from diverse data formats, including .fasta, .txt, .csv, .xlsx, and anndata files. The encoded representations can be seamlessly plugged into arbitrary perturbation effect prediction model —such as GEARS, CPA, or other models—to enable cross-domain perturbation modeling including G2CP.

## Author contributions

Y.L., J.Y., M.L. and F.Y. conceived the study. Y.L. developed the algorithm and conducted experiments under the supervision of F.Y. and J.Y.. L.L. and J.Z. provided valuable advice on UniPert framework and downstream task design. L.H. gave suggestions for the cross-domain transfer learning implications. Y.L. wrote the manuscript. Y.L. finished the figures under the guidance of F.Y. and J.Y.. M.Z. J.Y., F.Y., L.L, F.W., and M.L. revised the manuscript. J.Z. and F.Y. conducted the code review. All authors reviewed and approved the manuscript.

## Acknowledgements

The authors appreciate informative conversations with Yu Rong and Zetian Zheng, helpful visualization supports and suggestions from Shaoning Li, and Yin Fang. This work was partially supported by the National Natural Science Foundation of China (Grant 62225209 to M.L.), the Young Elite Scientists Sponsorship Program by CAST (2023QNRC001 to F.Y.), and the Fundamental Research Funds for the Central Universities of Central South University (No. 2023ZZTS0627 to Y.L.). We acknowledge Alex NG for his support in maintaining the AI for Life Sciences Lab. Extended Data Fig. 1a was created with BioRender.com.

## Extended Data Figures

**Extended Data Figure 1.**
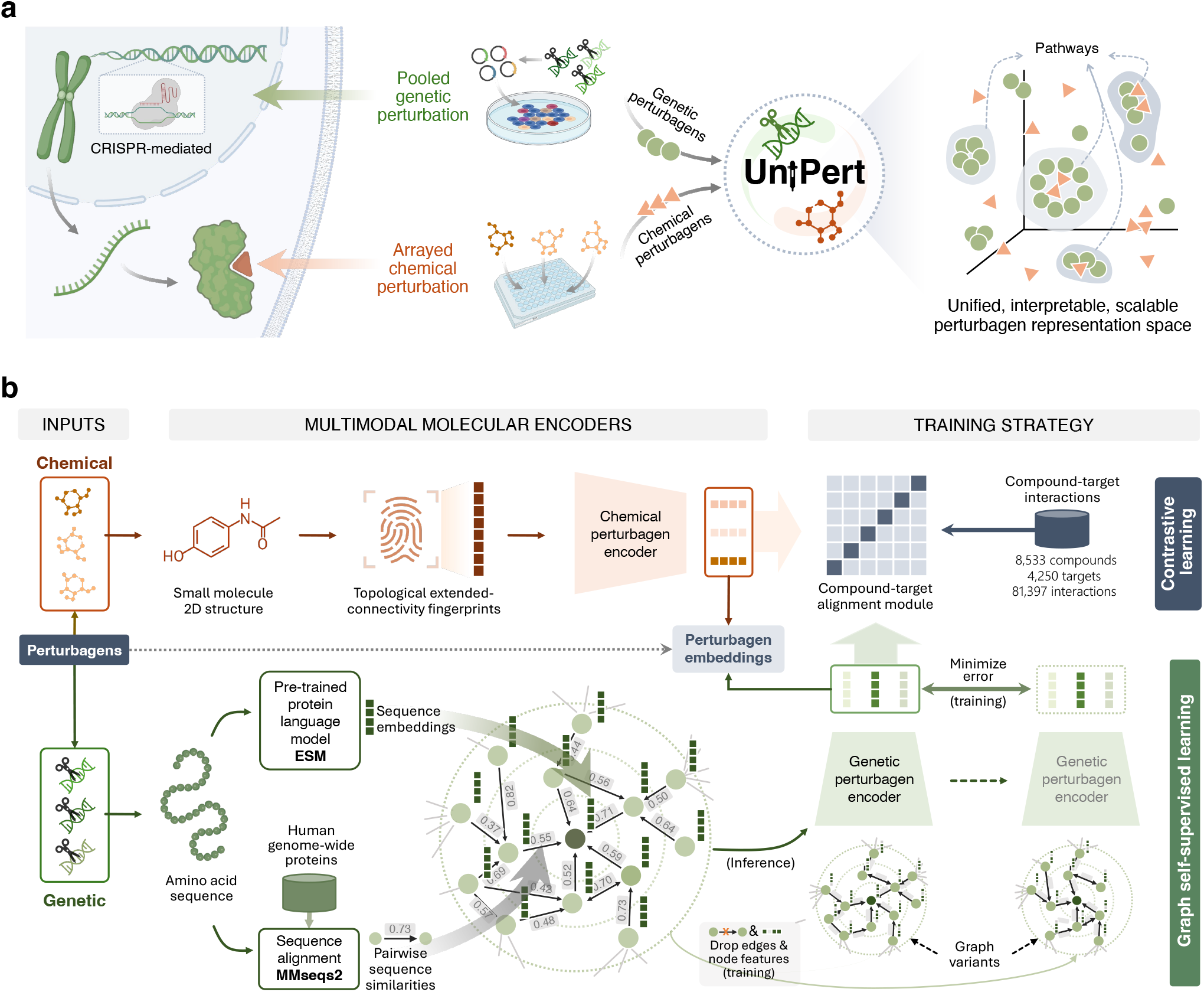
Illustration of UniPert model. **(a)** Problem formulation of UniPert: given genetic perturbagens (green) and chemical perturbagens (yellow), UniPert represents these large/small-scale molecules within a shared, biologically interpretable, and scalable representation space (right). **(b)** Hybrid design of UniPert: multi-scale molecular inputs (left), including amino acid sequences for genetic perturbagens and SMILES strings for chemical perturbagens, are processed through tailored model architectures (middle) trained by combined training strategies (right). For genetic perturbagens, the model combines sequence alignment algorithm, large PLM, and graph neural networks to preserve sequence features. For chemical perturbagens, binary ECFP fingerprints are adopted for capturing functional groups. Training involves graph self-supervised learning (bottom right) to refine genetic perturbagen embeddings and contrastive learning (top right) to align perturbagen modalities using experimentally validated compound-target interaction information.

**Extended Data Figure 2.**
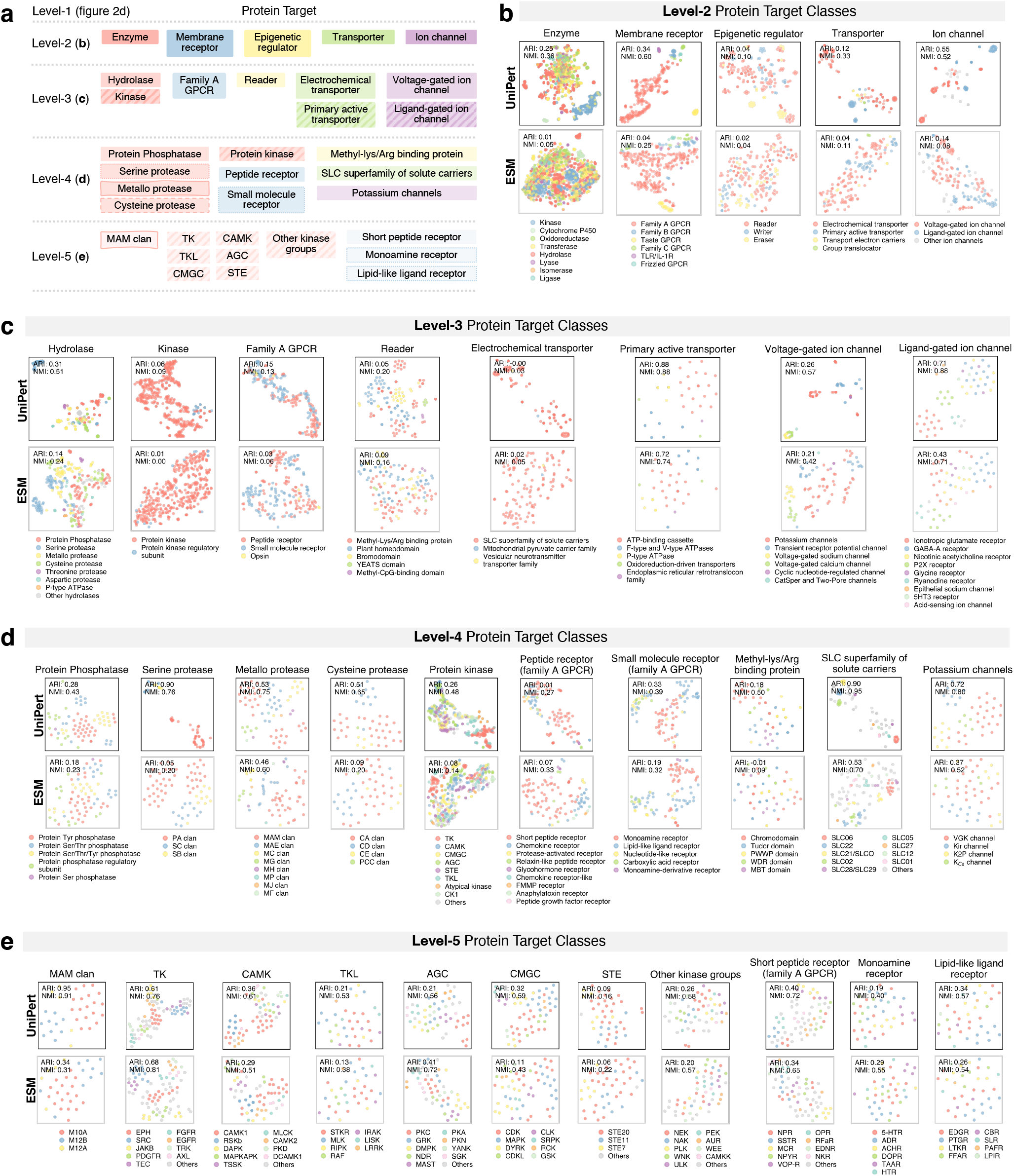
Performance comparison in clustering 35 hierarchical pharmacological target classes using UniPert and ESM embeddings. **(a)** Hierarchical relationships of protein target pharmacological classes, showing parent class names and relationships of classes (with more than 30 members) from level-2 to level-5. (**b-e)** t-SNE visualizations comparing UniPert (top panels, black-framed) and ESM (bottom panels, gray-framed) embeddings for clustering protein targets at different levels. Each panel represents to a specific target class at the corresponding pharmacological class level. Colors of scatters (represent proteins) correspond to distinct subclasses (legend) within each parent class (title). Clustering performance is evaluated using ARI and NMI.

**Extended Data Figure 3.**
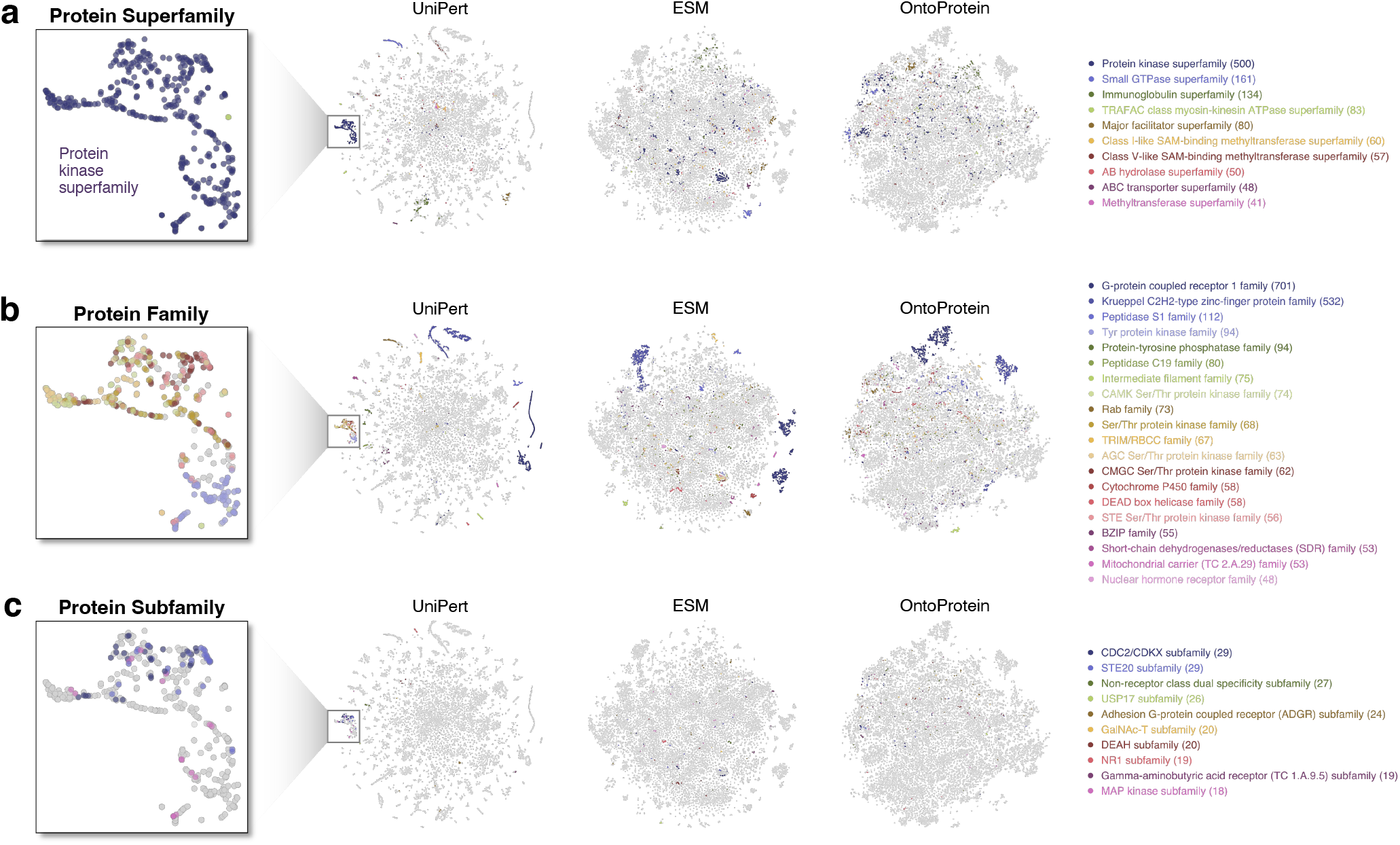
UniPert distinguishes hierarchical protein family classes in genome-wide protein embedding space. t-SNE visualizations of genome-wide protein (n = 19,443) using UniPert, along with state-of-the-art protein language models, ESM and OntoProtein. Colors denote classes of **(a)** the top 10 superfamilies, **(b)** the top 20 families, and **(c)** the top 10 subfamilies. The black frames (left) highlight the superior clustering capability of UniPert for the kinase superfamily.

**Extended Data Figure 4.**
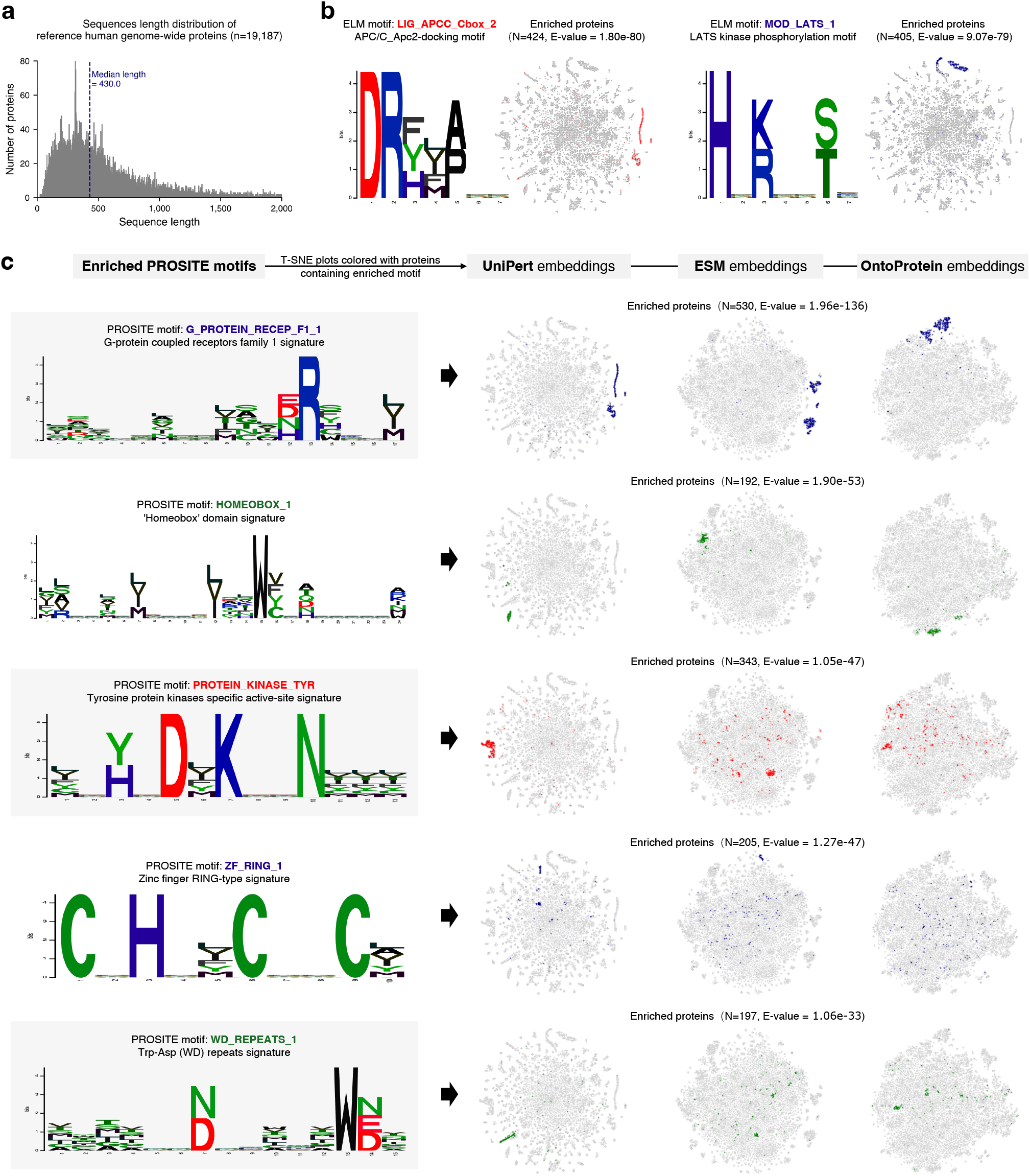
Functional motif enrichment analysis and clustering performance of reference genome-wide proteins. **(a)** Distribution of sequence lengths for reference human genome-wide proteins (n=19,187). The median sequence length is indicated. **(b)** Most significantly enriched motifs in the ELM motif database. Left: APC/C_Apc2-docking motif. Right: LATS kinase phosphorylation motif. **(c)** Most significantly enriched motifs in the PROSITE motif database. The E-value denotes the enrichment significance. t-SNE plots highlight proteins containing the motif in the genome-wide protein embedding space of UniPert, ESM, and OntoProtein.

**Extended Data Figure 5.**
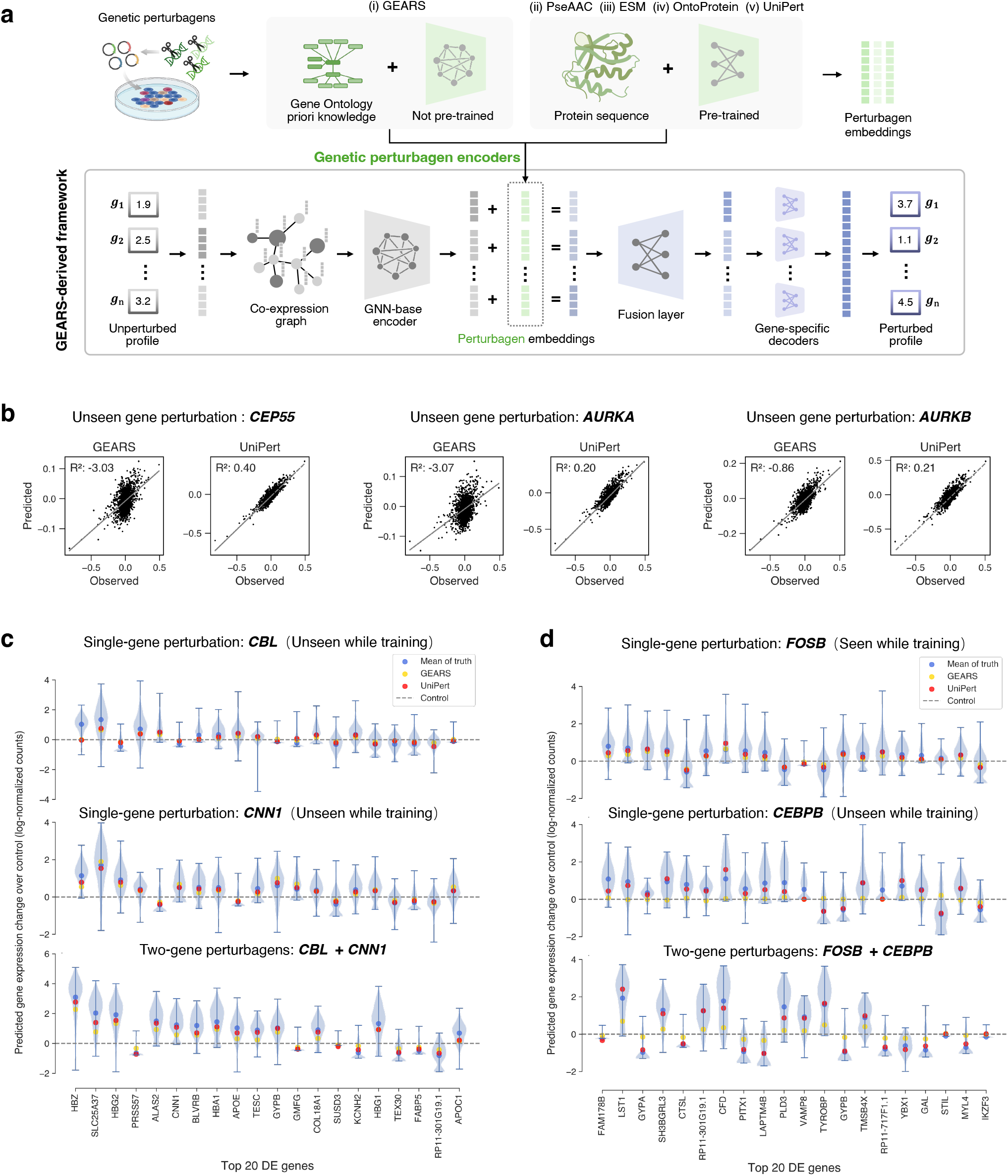
UniPert enhances unseen single and combination genetic perturbations effect prediction in the GEARS-derived framework. **(a)** Schematic of the GEARS-derived genetic perturbations effect prediction framework. Given unperturbed profiles (bottom left), i.e., the control group, each gene in the profile is encoded through GNNs with a gene co-expression graph. Embeddings of the genetic perturbagens (top left) can be learned by incorporating GO prior knowledge and GNNs, as in i) the original GEARS model, or extracted from extensible pre-trained/pre-defined protein sequence representation methods, such as ii) PseAAC, iii) ESM, iv) OntoProtein, and v) our proposed UniPert. Single/multiple perturbagen embeddings (green) are then added to each gene embedding (grey) to obtain post-perturbation gene embeddings (purple) that will be finally converted into predicted post-perturbation gene expression values after sequentially passing through the fusion and gene-specific decoders. **(b)** Scatterplots show real observed vs. predicted expression values using GEARS and UniPert after unseen single-gene perturbations (*CEP55, AURKA*, and *AURKB*) in the *Dixit et al*. dataset. **(c)** Case of the ‘seen 0’ scenario in combination gene perturbation prediction. Violin plots indicate distribution of real observed post-perturbation gene expressions values over control after unseen gene combination (*CBL*+*CNN1*) and unseen single-genes (*CBL* and *CNN1*) perturbation, and the colored markers show mean of real/predicted values. Top 20 DEGs of the combination perturbation in the *Norman et al*. dataset were displayed. **(d)** Case of the ‘seen 1’ scenario in combination gene perturbation prediction. This case shows comparison of unseen gene combination (*FOSB*+*CEBPB*), seen gene (*FOSB*), and unseen gene (*CEBPB*).

**Extended Data Figure 6.**
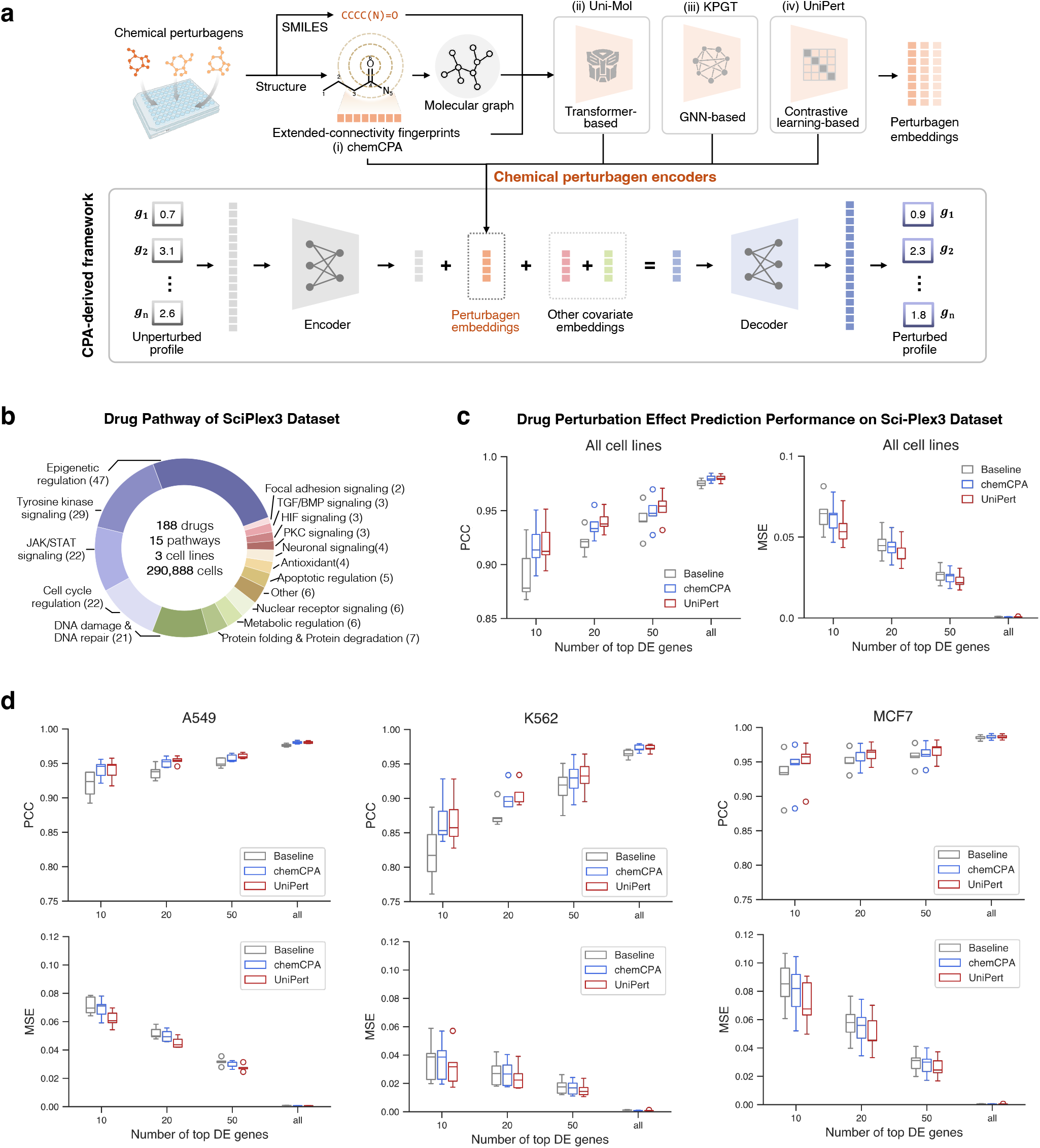
UniPert improves unseen chemical perturbation effect prediction performance in the CPA-derived framework. **(a)** Schematic of the CPA-derived chemical perturbations effect prediction framework. Unperturbed profile (bottom left) values are projected into a low-dimensional latent space, while chemical perturbagens (top left), i.e., small molecules, are numericized using traditional molecular fingerprint features like i) chemCPA model, or, be encoded with advanced representation methods such as ii) Uni-Mol, iii) KPGT, and our proposed iv) UniPert. Perturbagen embeddings (yellow) along with other covariate embeddings are then added to the latent control embedding (grey), producing predicted post-perturbation gene profiles (purple) after decoding. **(b)** Donut chart depicting the distribution of drug pathways in the large-scale drug screening dataset sciPlex3. **(c)** Overall performance comparison in predicting unseen drug perturbation effects using UniPert and chemCPA on the sciPlex3 dataset, evaluated at various DEG scales. **(d)** Performance comparison across three cell lines on the sciPlex3 dataset. The baseline represents real unperturbed profiles vs. real post-perturbed profiles

**Extended Data Figure 7.**
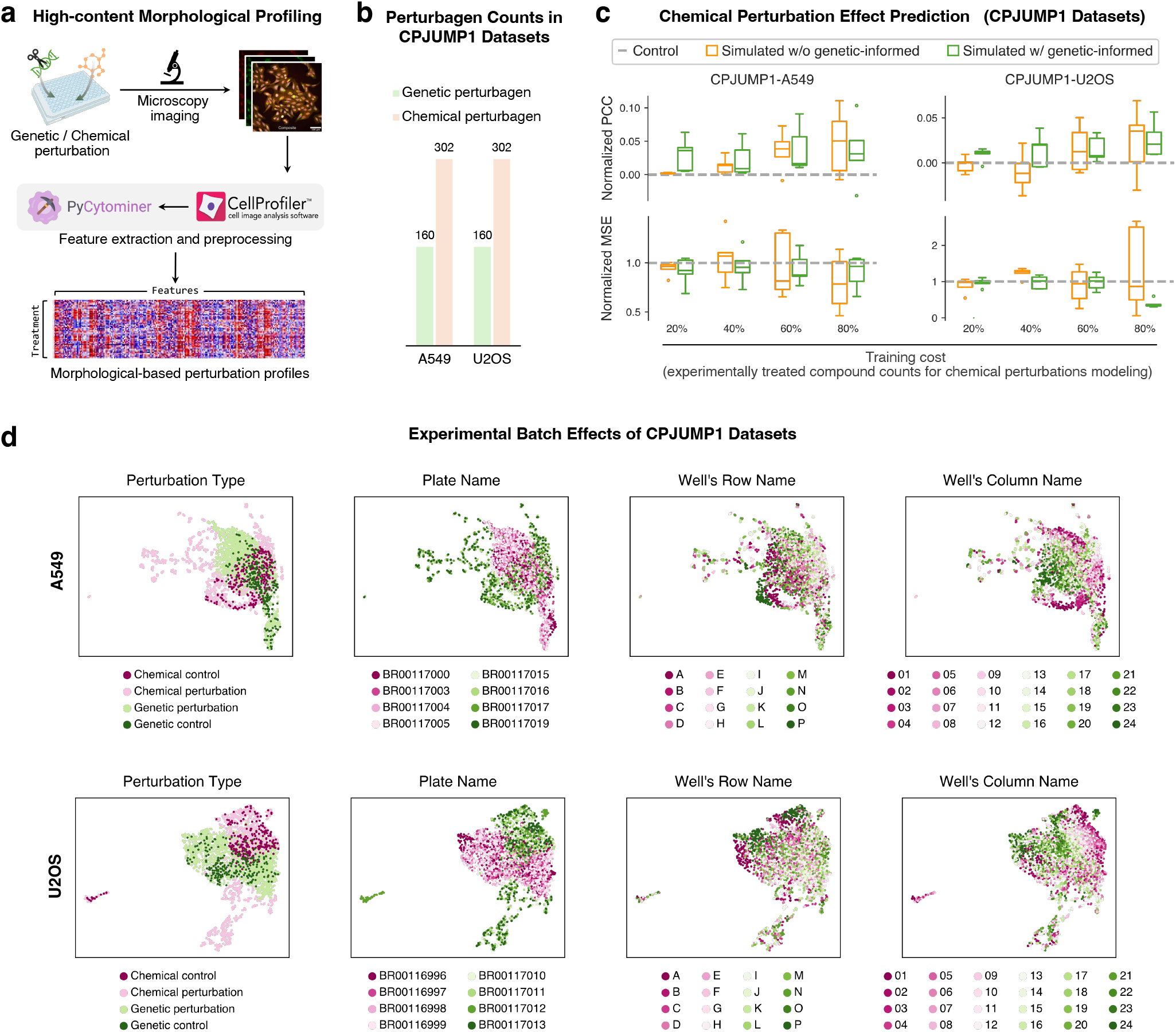
Application of G2CP to morphological perturbation profiles. **(a)** Workflow for generating morphological perturbation profiles. Genetic or chemical perturbation is followed by microscopy imaging and feature extraction using PyCytominer and CellProfiler, resulting in morphological-based profiles. **(b)** Perturbagen information of the CPJUMP1 datasets. Bar graphs show the number of genetic and chemical perturbagens in the two cell line datasets (A549 and U2OS). **(c)** Performance comparison in chemical perturbation effect prediction on the CPJUMP1 datasets. Box plots show the normalized PCC and normalized MSE resulting from G2CP (w/ genetic-informed) and prediction model without genetic perturbation pre-training (w/o genetic-informed). **(d)** Experimental batch effects of CPJUMP1 datasets. UMAP visualizations of CPJUMP1 datasets for A549 (top) and U2OS (bottom) cell lines, colored by perturbation type, plate name, well’s row name, and well’s column name, illustrating potential batch effects.

**Extended Data Figure 8.**
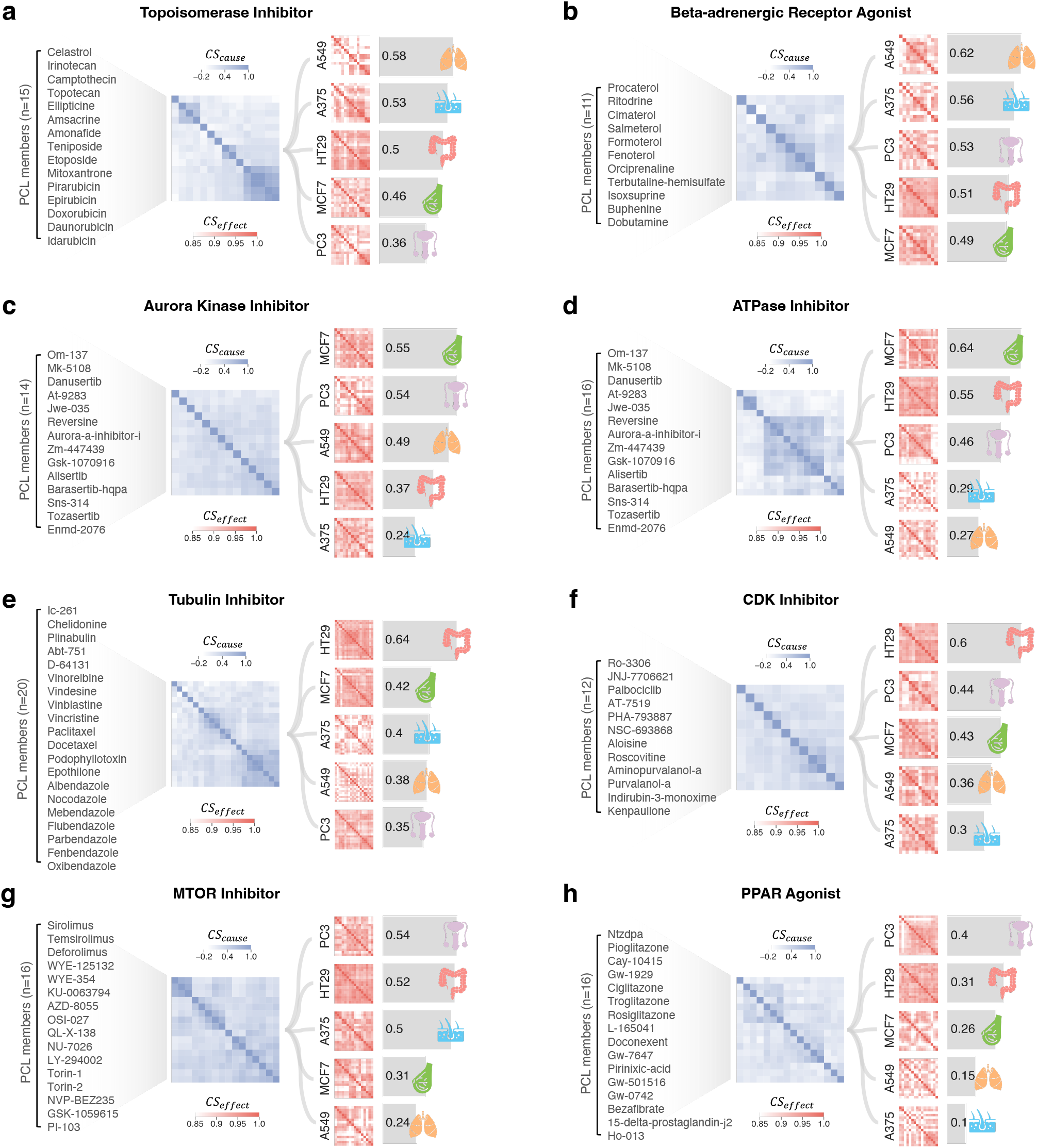
Charactered PCL cellular sensitivities across 5 cancer cell lines. Each panel illustrates the cellular sensitivities of the indicated PCL containing multiple drug members: **(a)** Topoisomerase inhibitor, **(b)** Beta-adrenergic receptor agonist, **(c)** Aurora kinase inhibitor, **(d)** ATPase inhibitor, **(e)** Tubulin inhibitor, **(f)** CDK inhibitor, **(g)** MTOR inhibitor, **(h)** PPAR receptor agonist.

